# Paclitaxel induces NM2-dependent cellular contraction through GEF-H1 dissociation from microtubules and RhoA/ROCK activation in cancer cells

**DOI:** 10.64898/2025.12.29.696873

**Authors:** Gloria Asensio-Juárez, Rafael Pérez-Díaz, Hugo Ramos-Solano, Marina Garrido-Casado, Vanessa C. Talayero, Miguel Vicente-Manzanares

**Affiliations:** Molecular Mechanisms Program, Centro de Investigación del Cáncer and Instituto de Biología Molecular y Celular del Cáncer, Consejo Superior de Investigaciones Científicas (CSIC)-Universidad de Salamanca, Spain

## Abstract

In this study, we have investigated the crosstalk between microtubule dynamics and actomyosin contractility in cancer cells treated with taxanes, which are chemotherapeutic agents used to treat solid tumors. We found that paclitaxel (PTXL) induced cell contraction through a mechanism that involved the rapid dissociation of GEF-H1 from microtubules, and the phosphorylation and acute activation of NM2 in a RhoA-dependent manner. Mutation of the major α-tubulin acetylation site, K40R, markedly slowed and reduced the efficiency of PTXL-induced GEF-H1 dissociation, indicating that K40 acetylation is required for the full, rapid release of GEF-H1 from the microtubule lattice. These findings were corroborated using inhibitors of tubulin deacetylase HDAC6, which promoted a slow release of GEF-H1 from microtubules and a lagged accumulation of phosphorylated NM2. Unexpectedly, depletion of tubulin acetyltransferase αTAT1 also induced NM2 phosphorylation, indicating that microtubule acetylation is involved in the maintenance of contractile homeostasis. Together, these results indicate that PTXL induces rapid cellular contraction dependent on the GEF-H1-RhoA-ROCK axis, in which K40 acetylation of α-tubulin gates the efficiency of GEF-H1 dissociation from microtubules, whereas homeostatic microtubule acetylation maintains appropriate levels of cellular contractility through long-term control of NM2 phosphorylation and actomyosin organization.

## 1. Introduction

Contractility is a major regulator of many cellular and tissue-level processes, including the homeostasis of epithelial monolayers, cell migration, wound healing, muscle contraction, cell differentiation and many other such processes (1). Contractility is mainly mediated by the action of myosin II molecular motors. Cells and tissues of various lineages and anatomical origins have a specific complement of myosin II paralogs that optimize their contractile functional output (2). For example, muscles mainly express muscle-specific myosin II variants, which are designed to form long and stable sarcomeres and produce very large, macroscopic forces by acting in unison. Conversely, non-muscle cells express non-muscle myosin II (NM2) paralogs, which form short and dynamic filaments that produce small bursts of force to drive local processes such as cell migration, cell division and most phenomena involving movement and/or resistance to deformation.

The crosstalk of class II myosins with cytoskeletal systems other than microfilaments is poorly documented. However, specific manipulations of microtubules have been shown to modulate contractility. For example, short-term (<3h) treatment with nocodazole increases cell contraction (3). Nocodazole is a cell-cycle synchronizing drug that is often used to arrest cells at prometaphase (4). It reversibly promotes microtubule depolymerization, preventing formation of the mitotic spindle that segregates sister chromatids between daughter cells. Interestingly, the short-term effect of nocodazole on contractility seems independent of its synchronizing capability. Instead, nocodazole promotes the release of GEF-H1, which is a potent RhoA activator (5), from the microtubule lattice into the cytoplasm, where it readily activates RhoA. GTP-bound (active) RhoA promotes cell contraction by activating NM2 (reviewed in (2)). The mechanism involves the activation of the Ser/Thr kinase ROCK, which directly phosphorylates Ser19 of the regulatory light chain of non-muscle and smooth muscle myosin II, promoting its conformational extension and activation (6, 7). ROCK also phosphorylates and inactivates the myosin II phosphatase MYPT1, which dephosphorylates Ser19 (8).

A similar effect to that of nocodazole on cellular contraction has been reported for other microtubule-depolymerizing drugs, such as colchicine (9) and vinblastine (VBL, this report). By binding to other sites of the microtubule structure (10), these compounds also trigger depolymerization, consistent with their similar effect on contractility. These two compounds are used clinically to treat gout, a local inflammatory reaction (reviewed in (11)), and various forms of solid cancer (reviewed in (11, 12), respectively).

Paclitaxel (PTXL) and docetaxel (DTXL) belong to the taxane family of chemotherapeutic drugs, which are used to treat a wide array of malignancies; like nocodazole, colchicine and VBL, taxanes also arrest cancer cells in prometaphase. Long-term arrest of cancer cells triggers cell apoptosis (13, 14). Their mechanism, however, is the reverse: PTXL and DTXL stabilize the microtubule lattice instead of depolymerizing it, blocking spindle reorganization (15), but still driving cells to mitotic arrest and apoptosis. In addition, while the activation of contractility by tubulin depolymerization drugs is well-characterized (5, 16–19), the effect of tubulin stabilizers on actomyosin dynamics is unclear. This is of critical importance due to the widespread use of PTXL and DTXL in the treatment of solid tumors (20). In this project, we sought to characterize the effect of taxane-induced tubulin stabilization on actin dynamics, particularly through its potential impact on NM2 activity. We found that PTXL also induced NM2 activation and an overall increase in contractility. Mechanistic observations revealed that PTXL has a dual positive effect on NM2 activation: similar to tubulin depolymerizers, it promotes GEF-H1 dissociation from the microtubule lattice, increasing NM2 phosphorylation and overall contractility throughout the cell. Importantly, the kinetics of GEF-H1 dissociation from microtubules induced by PTXL were much faster than those induced by inhibitors of HDAC6 (21), which promote GEF-H1 release in a much slower manner. A mutation of the major αTAT1-dependent acetylation site in α-tubulin, K40R, partially inhibited PTXL-induced GEF-H1 release from microtubules and completely countered the effect of tubacin, a HDAC6 inhibitor. Interestingly, depletion of the tubulin acetyltransferase αTAT1 (22, 23) did not decrease total RLC phosphorylation, but increased its accumulation in the lamella, concomitant with impaired anteroposterior cell polarization. Finally, PTXL did not promote the association of UNC45a with NM2, an interaction that is a mandatory step of filament assembly (24), indicating that the main mechanism of increased contractility involves the GEF-H1/RhoA/ROCK-dependent activation of NM2 and not enhanced UNC45a-dependent filament assembly. Together, our data indicate that PTXL increases contractility by promoting RLC phosphorylation in a GEF-H1/RhoA-dependent manner similar to that of microtubule polymerization inhibitors. They also suggest that microtubule-dependent stimulation of contractility follows two possible routes: i) a tonic increase in NM2 activation triggered by the suppression of microtubule acetylation; ii) the acute phosphorylation of NM2 by treatment with PTXL or microtubule polymerization inhibitors in a GEF-H1/RhoA-dependent manner. We speculate that such mechanisms may be part of the development of resistance (25–27), which may lead to enhanced metastasis by increasingly motile cells.

## 2. Results

### 2.1. PTXL increases NM2 phosphorylation at Ser19 and Thr18/Ser19 and promotes cellular contraction

Previous studies have shown that chemically induced microtubule depolymerization, e.g. treatment with nocodazole, increases NM2 phosphorylation at Ser19 of the RLC (5). Taxanes have the opposite effect on microtubules to that of nocodazole, that is, they induce microtubule stabilization and inhibit depolymerization (15). We thus hypothesized that taxanes could decrease NM2 phosphorylation. To test this hypothesis, we treated U2OS osteosarcoma cells with 100 nM PTXL for 3h. These conditions promoted rapid accumulation of acetylated microtubules, as revealed by Western blot using a specific antibody **(Fig. 1A)**. As a control, we used VBL, which is a microtubule depolymerizing drug that has been used in tumor chemotherapy (12). Similar to nocodazole (28), VBL induced tubulin depolymerization, significantly reducing the amount of acetylated tubulin **(Fig. 1A)**. It also increased RLC phosphorylation at Ser19 **(Fig. 1B-C)**. Strikingly, PTXL also promoted RLC phosphorylation at Ser19 by Western blot **(Fig. 1B-C)** and immunofluorescence **(Fig. 2C)**. Similar results were also observed in two ovarian cancer cell lines, OVCAR-8 and SK-OV-3 **(Fig. 1D-G)**. PTXL also increased cell contraction, as revealed by a significant decrease in the spreading area of osteosarcoma cells on fibronectin **(Fig. 2A-B)**. In treated cells, we observed an increase in both phospho-Ser19 and phospho-Thr18/Ser19 RLC **(Fig. 1B-C)**, and the appearance of elongated fibers decorated with phospho-Ser19 RLC, which were seldom observed in control cells **(Fig. 2C, insert)**. It also increased the number and size of focal adhesions in cells spread on fibronectin **(Fig. 2E-G)**. Adhesions were significantly larger, as illustrated by their perimeter, and this correlated with increased adhesion length (**Fig. 2D**, insert and **Fig. 2H**). This is a typical feature of cells displaying increased contractility (29). Decreased cell area and increased adhesion elongation together with the elevation of phospho-Ser19 RLC levels are clear indicators of increased contractility (29, 30). These results indicate that microtubule depolymerizing agents and microtubule stabilizers, despite having antagonistic effects on microtubule stability, produce comparable effects on the contractile machinery of adherent tumor cells.

**Figure 1:**
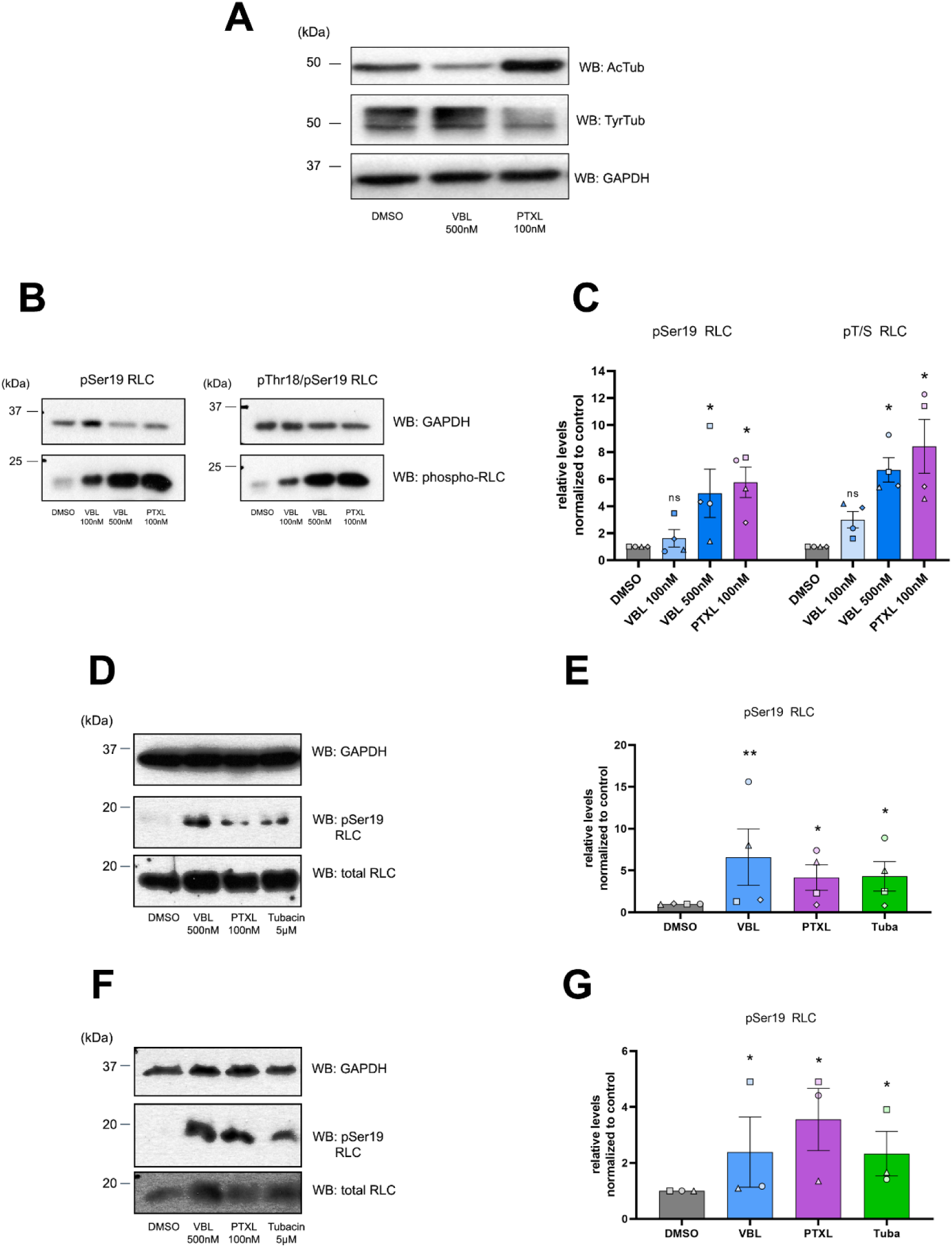
Paclitaxel increases NM2 phosphorylation in osteosarcoma and ovarian cancer cells. (A) U2OS osteosarcoma cells were treated with VBL or PTXL (or DMSO) drugs for 3 hours, lysed, proteins separated by SDS/PAGE and blotted for acetylated and tyrosinated tubulin. GAPDH is shown as loading control. (B) U2OS cells were treated with VBL (100 nM or 500 nM) or PTXL (100 nM) for 3 hours, lysed, proteins separated by SDS/PAGE and blotted for phospho-Ser19 RLC, phospho-Thr18/Ser19 RLC and GAPDH, shown as loading control. A representative experiment of five performed is shown. (C) Quantification of the phosphorylation levels shown in (B). Data are the mean ± SD of five independent experiments. (D-E) OVCAR-8 ovarian cancer cells were treated with the indicated inhibitors for 3h, lysed, proteins separated by SDS/PAGE and blotted as indicated. A representative Western blot experiment of four performed is shown. Quantification of the four experiments is shown in (E). ** p<0.01; * p<0.05 (Mann-Whitney U test). (F-G) SK-OV-3 ovarian cancer cells were treated with the indicated inhibitors for 3h, lysed, proteins separated by SDS/PAGE and blotted as indicated. A representative Western blot experiment of three performed is shown. Quantification of the three experiments is shown in (G). * p<0.05 (Mann-Whitney U test).

**Figure 2.**
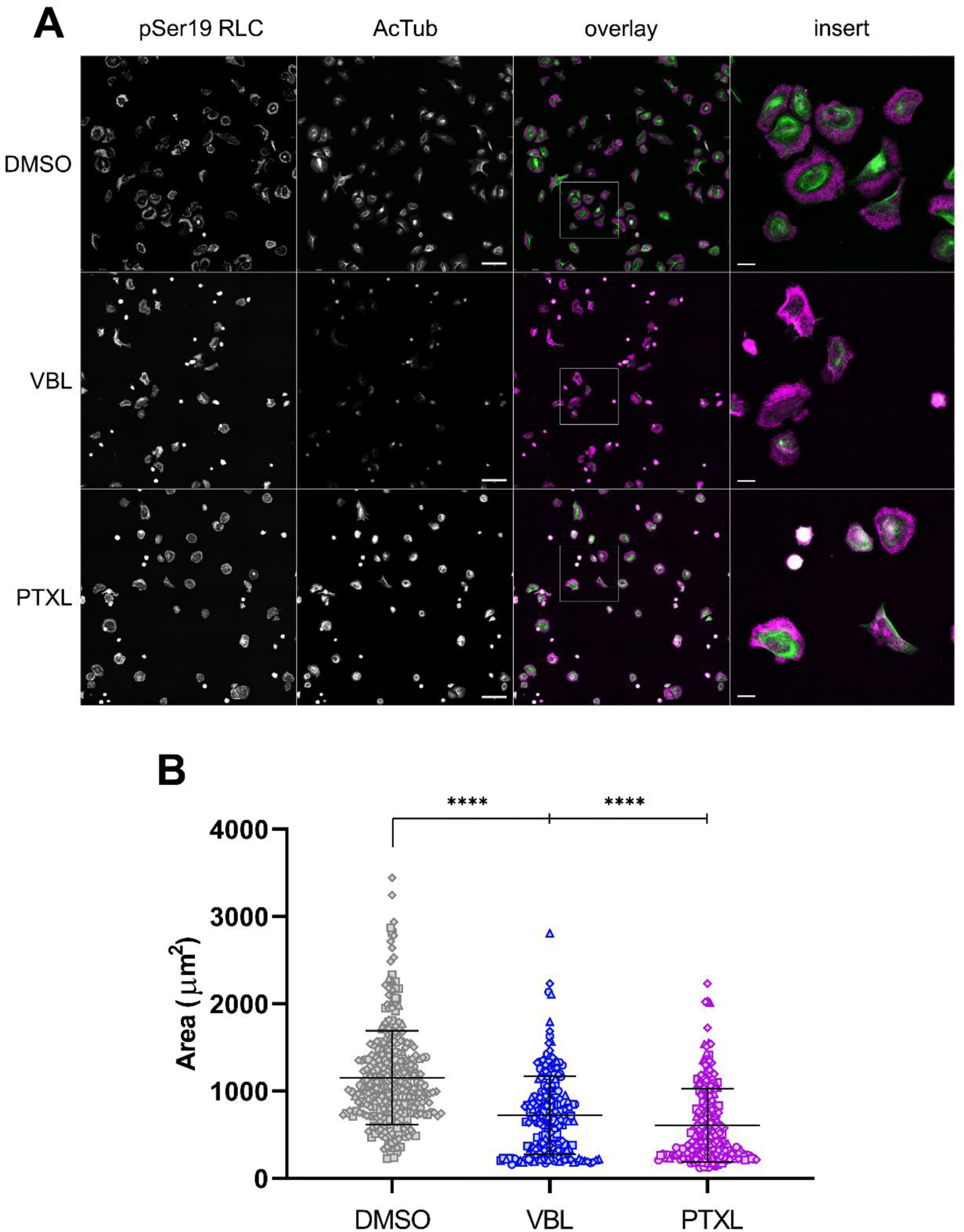
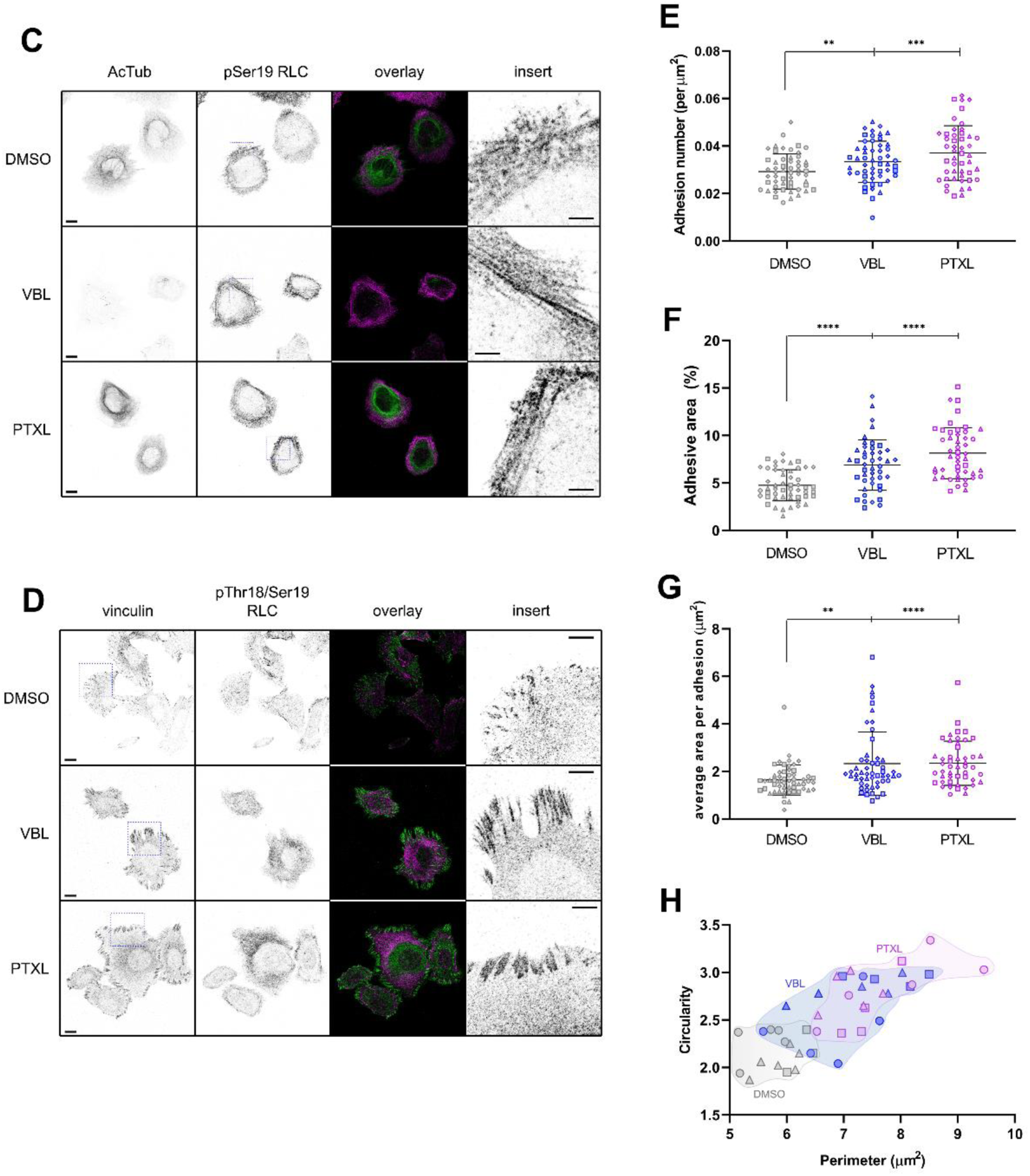
Paclitaxel increases cell contraction and promotes adhesion maturation and elongation. (A) Representative images of U2OS cells spread on 2 µg/mL of fibronectin overnight, then treated with anti-mitotic drugs for 3h, fixed, permeabilized and stained as indicated. Green represents acetylated tubulin and red represents phospho-Ser19 RLC. Scale bars = 50 µm, large image; =10 µm, insert. (B) Quantification of cellular area as shown in (A). Data correspond to >200 cells from four independent experiments. Every experiment is pooled in the graph, and symbols denote data from different replicates. Significance was determined using non-parametric Mann-Whitney U test. **** p<0.0001. (C) Representative fluorescence images of U2OS cells stained for acetylated tubulin (AcTub) and phospho-Ser19 RLC. Scale bars = 10 µm, large image; =5 µm, insert. Inserts correspond to the squared regions (indicated by dotted lines) shown in the second column and represent endogenous pSer19 RLC. (D) Representative fluorescence images of U2OS cells stained for vinculin and phospho-Thr18/Ser19 RLC. Scale bars = 10 µm, large image; =5 µm, insert. Inserts correspond to the squared regions (indicated by dotted lines) shown in the first column and represent endogenous vinculin. (E-H) Quantification of the number of adhesions per μm^2^ (E); percentage of adhesive area (F), representing the area of the cell positive for vinculin divided by the total area of the cell; (G) average area per adhesion; and (H) adhesion perimeter with respect to adhesion elliptical index (circularity), defined by vinculin staining as calculated in (Talayero and Vicente-Manzanares, 2020). Each data point represents a group of n≥20 cells from four independent experiments, each designated by a different symbol. Significances were determined using Mann-Whitney U test. ** p<0.01; *** p<0.001; **** p<0.0001.

### 2.2. PTXL-induced NM2 phosphorylation depends on the RhoGEF GEF-H1

Previous reports demonstrated that nocodazole activates NM2 in a RhoA-dependent manner by activating a microtubule-associated RhoGEF, GEF-H1 (ARHGEF2) (5). GEF-H1 increases RLC pSer19 by activating the RhoA/ROCK axis, which phosphorylates and inactivates the myosin-specific phosphatase MYPT1 (8). To study whether the observed effect of PTXL on phosphorylation of RLC at Ser19 was dependent on GEF-H1, we generated GEF-H1-depleted U2OS cells using a specific shRNA sequence (31). Efficient silencing in U2OS cells was verified by flow cytometry and Western blot **(Fig. 3A-B)**. Next, we treated GEF-H1-depleted or control cells with 100 nM PTXL for 3h. GEF-H1 silencing impaired VBL- and PTXL-induced phosphorylation at Ser19 and Thr18/Ser19 **(Fig. 3B-C)**. Interestingly, GEF-H1 depletion decreased the levels of NM2-B heavy chain (*Myh10)*, but not of NM2-A heavy chain (*Myh9)* **(Fig. 3D-E)**, suggesting that different feedback regulatory circuits control the expression of each isoform. GEF-H1 depletion decreased NM2-B levels similarly across DMSO, VBL, and PTXL treatment conditions, consistent with an effect on NM2-B expression rather than a treatment-specific response **(Fig. 3E)**.

**Figure 3.**
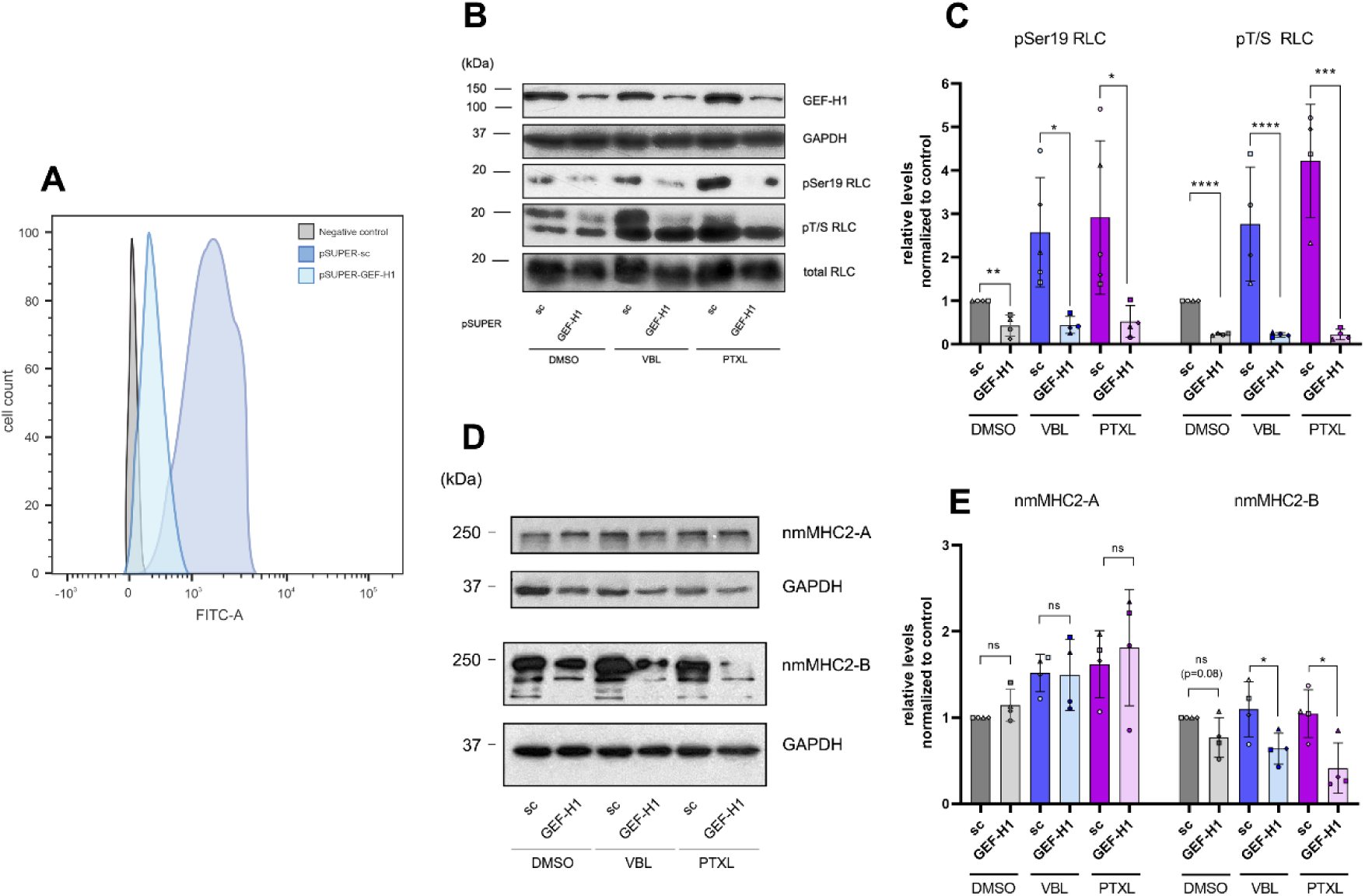
Microtubule poisoning-dependent phosphorylation of RLC at Ser19 requires normal GEF-H1 levels. (A) Representative experiment showing the reduction of GEF-H1 expression by pSUPER-GEF-H1 measured by flow cytometry in U2OS cells after 72h transfection. Control cells were transfected with pSUPER carrying a control RNA sequence (scrambled). Experiment is representative of eight performed. (B) Expression of GEF-H1 and phosphorylation of RLC at Ser19 and Thr18/Ser19 in GEF-H1-silenced U2OS cells treated with VBL or PTXL. Total RLC and GAPDH are shown as loading controls. A representative experiment of three performed is shown. (C) Quantification of phosphorylation protein levels from silenced cells as in (B). Data are the mean ± SD of four independent experiments. Significance was determined as indicated using Mann-Whitney U test. * p<0.05; ** p<0.01; *** p<0.001; **** p<0.0001. (D) GEF-H1-silenced U2OS cells were treated with VBL or PTXL for 3 hours, lysed, proteins separated by PAGE/SDS and blotted against nmMHC2-A and nmMHC2-B. GAPDH is shown as loading control. A representative experiment of three performed is shown. (E) Quantification of NM2 paralogs levels from silenced cells as in (D). Data are the mean ± SD of four independent experiments. Significance was determined as indicated using Mann-Whitney U test. ns, not significant; * p<0.05; ** p<0.01; *** p<0.001; **** p<0.0001.

Consistent with its effect on RLC phosphorylation, GEF-H1 depletion abrogated the VBL- and PTXL-induced increase in focal adhesion number and size seen in scrambled shRNA controls **(Fig. 4A-B, D-F)**. It also prevented the decrease in cellular area induced by VBL and PTXL **(Fig. 4C)**. shRNA-mediated depletion of RhoA, the downstream target of GEF-H1, similarly impaired VBL- and PTXL-induced RLC phosphorylation **(Fig. 5A-B).** Together, these results indicate that, similar to microtubule polymerization inhibitors, taxanes increase contractility through a GEF-H1/RhoA-dependent pathway.

**Figure 4.**
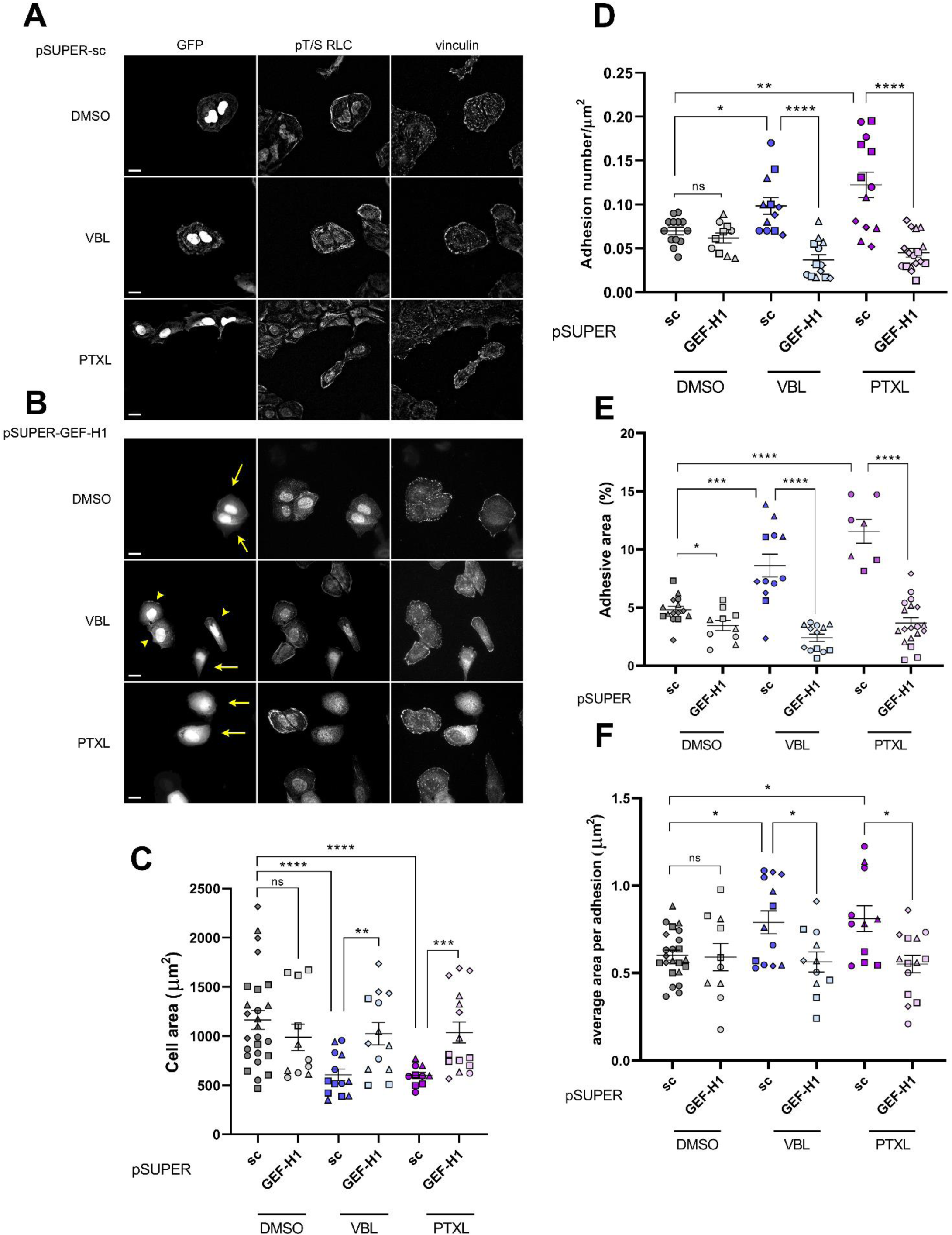
The effect of PTXL on cell size and adhesion morphology requires normal levels of GEF-H1. (A-B) Representative fluorescence images from U2OS control (A) or GEF-H1-silenced (B) cells treated with anti-mitotic drugs and spread on fibronectin 2 µg/mL coverslips. LifeAct-GFP was co-transfected in a 1:10 ratio to identify cells that had incorporated the depletion plasmid. We also visualized phospho-Thr18/Ser19 RLC and endogenous vinculin. Arrows indicate fully silenced cells, and arrowheads point to cells displaying partial silencing. Note the disappearance (arrows) and decrease (arrowheads) in pT/S RLC staining in bundles. Nuclear intensities denote non-specific binding. Bars = 20 µm. (C) Quantification of total cell area, determined by the perimeter of the cell as visualized by LifeAct-GFP. Data are the mean ± SD of four independent experiments, and data points represent >50 cells per experiment, each identified by a different symbol. (D-F) Quantification of the number of adhesions per μm^2^ (D); adhesive area (E) and average area per adhesion (F). Data are the mean ± SD of four independent experiments, and data points represent groups of 8-10 cells, each identified by a different symbol. Significances were determined using Mann-Whitney U test. ns, not significant; * p<0.05; ** p<0.01; *** p<0.001; **** p<0.0001.

**Figure 5.**
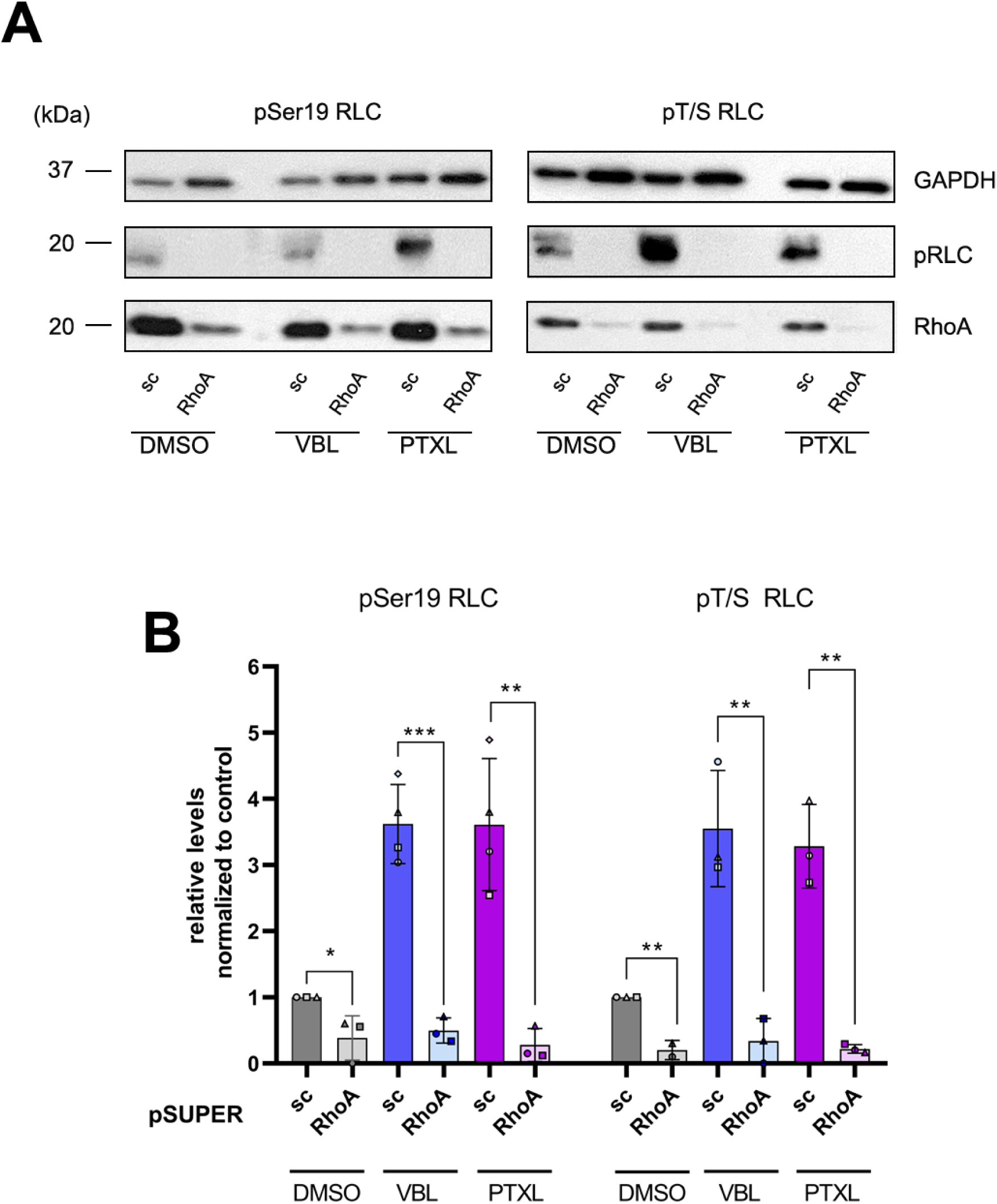
RLC phosphorylation at Ser19 and Thr18/Ser19 by PTXL requires normal levels of RhoA. (A) Control- or RhoA-silenced U2OS cells were treated with VBL or PTXL for 3 hours, lysed and blotted for phospho-Ser19, phospho-Thr18/Ser19 RLC and RhoA. GAPDH is shown as loading control. A representative experiment of three performed is shown. (B) Quantification of RLC phosphorylation from RhoA-silenced cells as in (A). Data are the mean ± SD of four independent experiments. Significance was determined with respect to DMSO-treated cells using Mann-Whitney U test. * p<0.05; ** p<0.01; *** p<0.001.

### 2.3. PTXL decreases the interaction of GEF-H1 with microtubules

GEF-H1 activation by microtubule polymerization inhibitors involves depolymerization of the microtubule lattice, releasing GEF-H1 into the cytoplasm in its free, unbound form. GEF-H1 then activates membrane-bound RhoA (32), triggering downstream signals that elevate cellular contractility in an NM2-dependent manner. To test whether the effect of PTXL on NM2 also relies on the dissociation of GEF-H1 from microtubules, we examined whether VBL and PTXL affected the co-sedimentation of GEF-H1 with microtubules. We found that VBL and PTXL had a similar inhibitory effect on the co-sedimentation of GEF-H1 with microtubules, displacing a significant portion of GEF-H1 to the supernatant **(Fig. 6A-B)**. Importantly, addition of VBL or PTXL to the fractionation buffer increased the extent of the effect **(Fig. 6A-B)**. To further probe the localization of GEF-H1 in cells treated with PTXL and VBL, we examined the colocalization of GFP-GEF-H1 with mCherry-α-tubulin. We found that GFP-GEF-H1 colocalized with the mCherry-decorated microtubule lattice **(Fig. 7A-B)**. Conversely, GFP-GEF-H1 rapidly diffused in the cytoplasm upon treatment with either PTXL or VBL **(Fig. 7A-D)**. These data indicate that PTXL also decreases the interaction of GEF-H1 with microtubules. Time-lapse analysis using TIRF microscopy revealed that VBL- and PTXL-induced dissociation of GFP-GEF-H1 from microtubules was almost instantaneous **(Fig. 7A-D)**.

**Figure 6.**
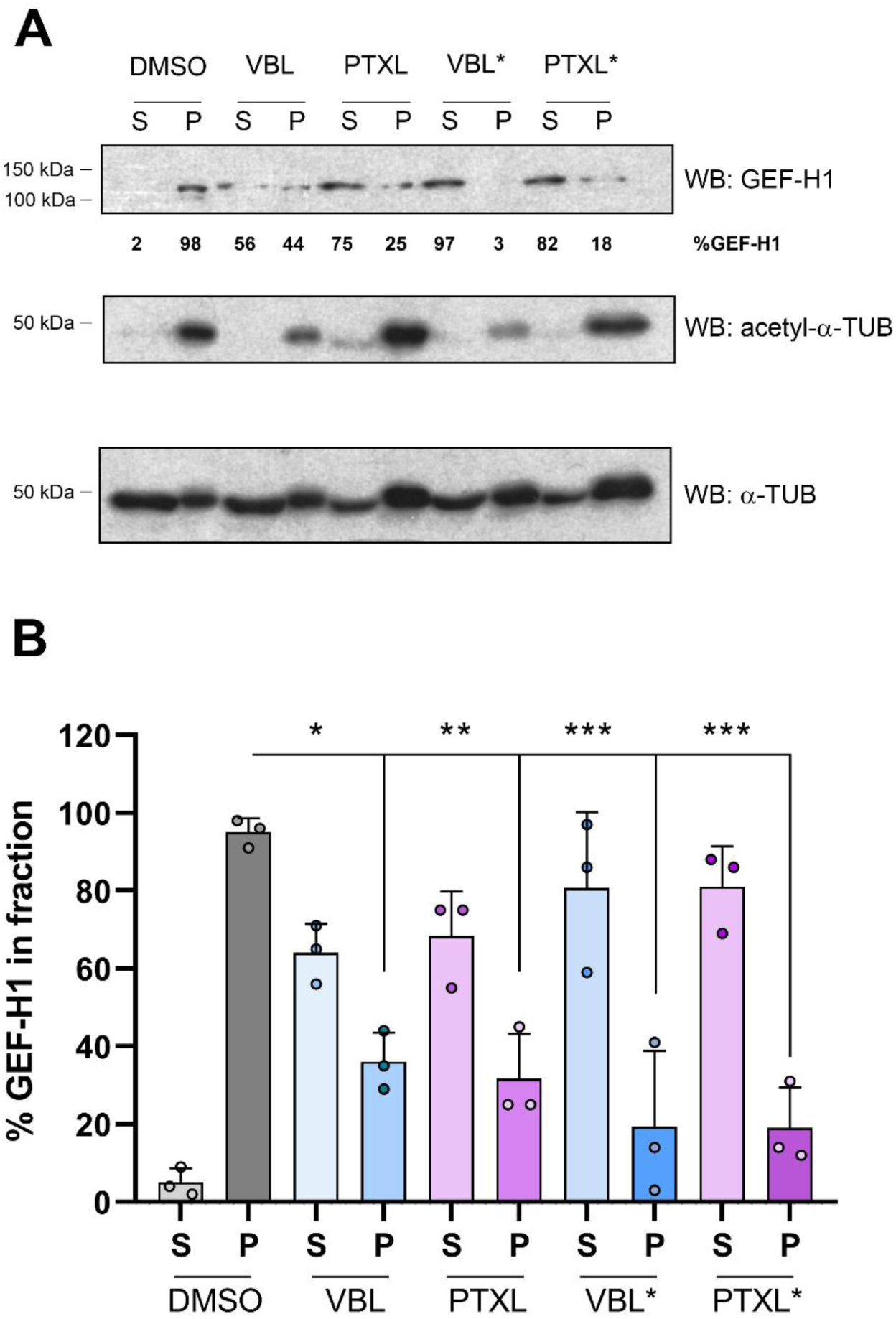
PTXL decreases the co-sedimentation of GEF-H1 with microtubules. (A) NIH-3T3 cells were treated with VBL or PTXL for 3h and fractionated as indicated in Materials and Methods to separate tubulin monomers/dimers (S = supernatant) from tubulin polymers (P = pellet). VBL* and PTXL* denote conditions in which the inhibitor was also included in the lysis buffer. Acetylated tubulin is also shown as a control for inhibitor activity and total tubulin as a marker of fractionation. The number below each band corresponds to the relative amount of GEF-H1 present in each fraction. Experiment is representative of three performed. (B) Quantification of GEF-H1 fractionation as in (A). Data are the mean ± SD of three independent experiments. Significance was determined with respect to DMSO-treated cells using Mann-Whitney U test. * p<0.05; ** p<0.01; *** p<0.001.

**Figure 7.**
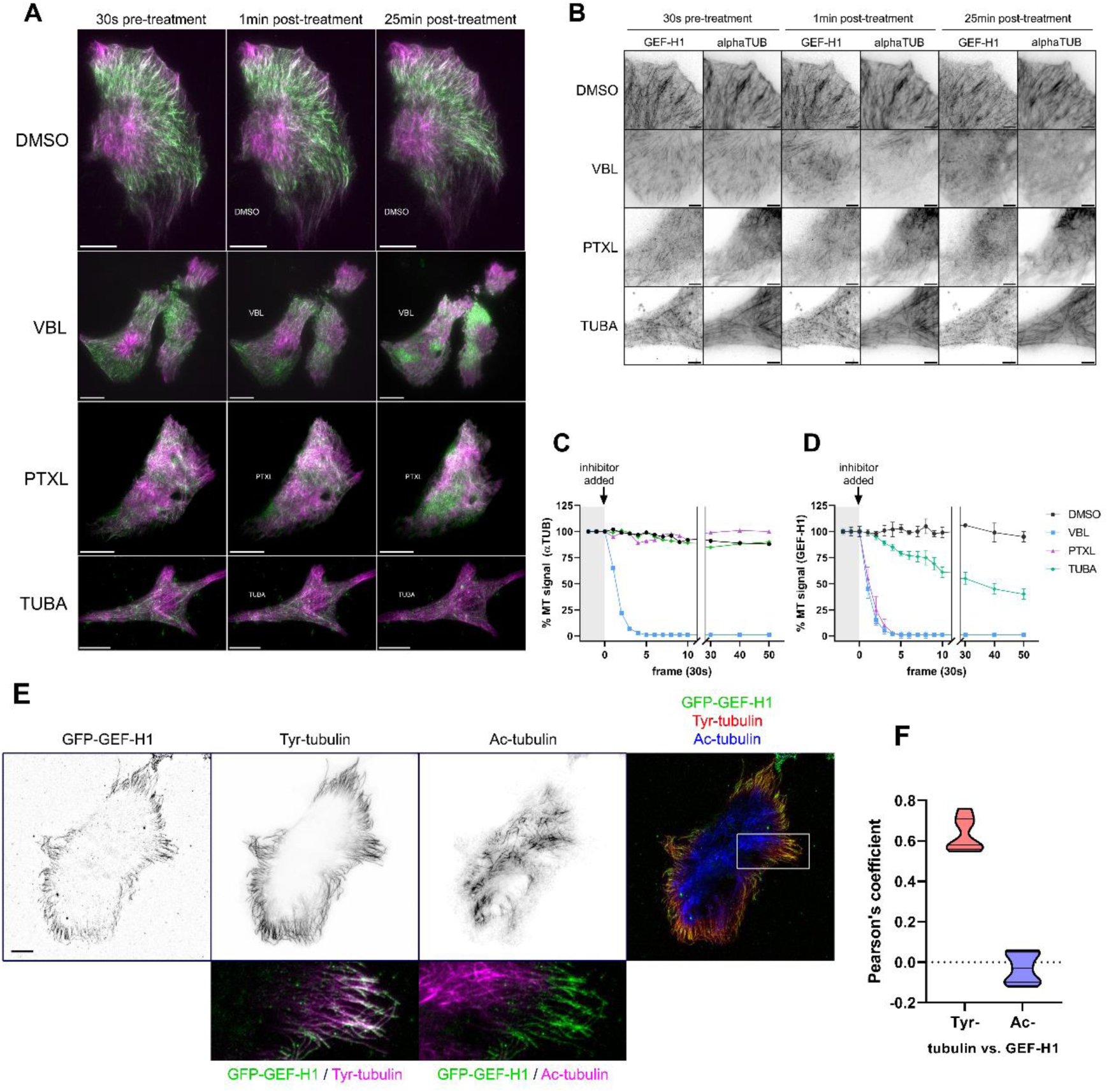
PTXL promotes the release of GEF-H1 from microtubules. (A) Representative TIRF microscopy images from time-lapse movies of U2OS cells co-transfected with mCherry-α-tubulin and GFP-GEF-H1. After 72 hours of co-transfection, cells were allowed to spread on fibronectin 2µg/mL coverslips. The next day, cells were placed under the microscope and treated with DMSO, 500 nM VBL, 100 nM PTXL or 5 µM tubacin (TUBA) at time=0; then were imaged for two additional hours. Left column represents a frame just before treatment, middle column 1 min post-treatment and right column 25 min post-treatment. Note the rapid disassembly of microtubules and GEF-H1 in VBL-treated cells; the immediate disappearance of GEF-H1 from condensing microtubular structures in PTXL-treated cells; and the lagged decrease in fluorescence in the case of tubacin-treated cells. Bars = 20 µm. Images taken from supplementary videos 1-4. (B) Black and white detail of the panels shown in (A) that illustrate more clearly the distribution of GFP-GEF-H1 and mCherry-α-tubulin at the indicated time points. Note the dissipation of GFP-GEF-H1 and mCherry-α-tubulin in the VBL panel at 25 min; the coalescence of mCherry-α-tubulin on the top-right corner of the image in the PTXL-treated cell, whereas GFP-GEF-H1 no longer follows a microtubular pattern; and the modest effect of TUBA in both markers at this time point. (C-D) Representative kinetics of MT signal of mCherry-α-tubulin (C) and GFP-GEF-H1 (D). Quantification was performed as indicated in Material and Methods out of 12 independent movies per condition from three independent experiments. (E) Representative TIRF microscopy images from fixed U2OS cells on fibronectin, expressing GFP-GEF-H1 and stained for tyrosinated (Tyr-) and acetylated (Ac-) tubulin. Lower panels illustrate the colocalization of GFP-GEF-H1 with tyrosinated microtubules and its segregation from acetylated microtubules. (F) Quantification of colocalization expressed as Pearson’s correlation coefficient (PCC), calculated per cell using the Coloc2 plugin (Fiji/ImageJ) after background subtraction and Costes automatic thresholding. n = 10-20 cells per condition, pooled from three independent experiments. p < 0.001 (Mann-Whitney U test).

Previous reports have indicated that tubulin acetylation is involved in the mechanosensitive response that triggers focal adhesion elongation and increased contractility via GEF-H1 release (21, 33). We thus hypothesized that treatment with HDAC6 inhibitors, which increase tubulin acetylation, would also promote GEF-H1 release from microtubules (21). To address whether microtubule acetylation similarly modulates the dissociation of GEF-H1 observed upon PTXL treatment, we examined the effect of tubacin, a specific HDAC6 inhibitor, on GEF-H1 localization. Time-lapse, live-image TIRF microscopy of cells co-expressing GFP-GEF-H1 and mCherry-α-tubulin revealed that tubacin induced a lagged dissociation of GEF-H1 from microtubules compared to the near-instantaneous effect of PTXL **(Fig. 7A-B, D)**. In addition, we observed a clear preference for GFP-GEF-H1 to interact with tyrosinated, dynamic microtubules over acetylated, stable microtubules **(Fig. 7E-F)**, consistent with acetylation status influencing the steady-state association of GEF-H1 with the microtubule lattice. These observations raised the question of whether K40, the major acetylation site in α-tubulin (34), controls the efficient dissociation of GEF-H1 induced by PTXL. To address this directly, we co-expressed GFP-GEF-H1 with a mCherry-tagged α-tubulin mutant (K40R) that cannot be acetylated at this site. Microtubules remained stable throughout the treatment with PTXL or tubacin **(Fig. 8A-B)**. PTXL induced a markedly slower and less complete dissociation of GEF-H1 from K40R microtubules **(Fig. 8A,C)** compared to the rapid, near-complete dissociation observed from wild-type microtubules in the same experiments **(Fig. 7A-B, D)**. Tubacin had no statistically significant effect on GEF-H1 association with K40R microtubules **(Fig. 8A,C)**, consistent with K40 acetylation being tubacin’s principal mode of action in this context. Together, these data indicate that K40 acetylation of α-tubulin is required for the efficient, rapid dissociation of GEF-H1 induced by PTXL, and that the residual, slower dissociation observed on K40R microtubules reflects a K40-independent component of the PTXL response.

**Figure 8.**
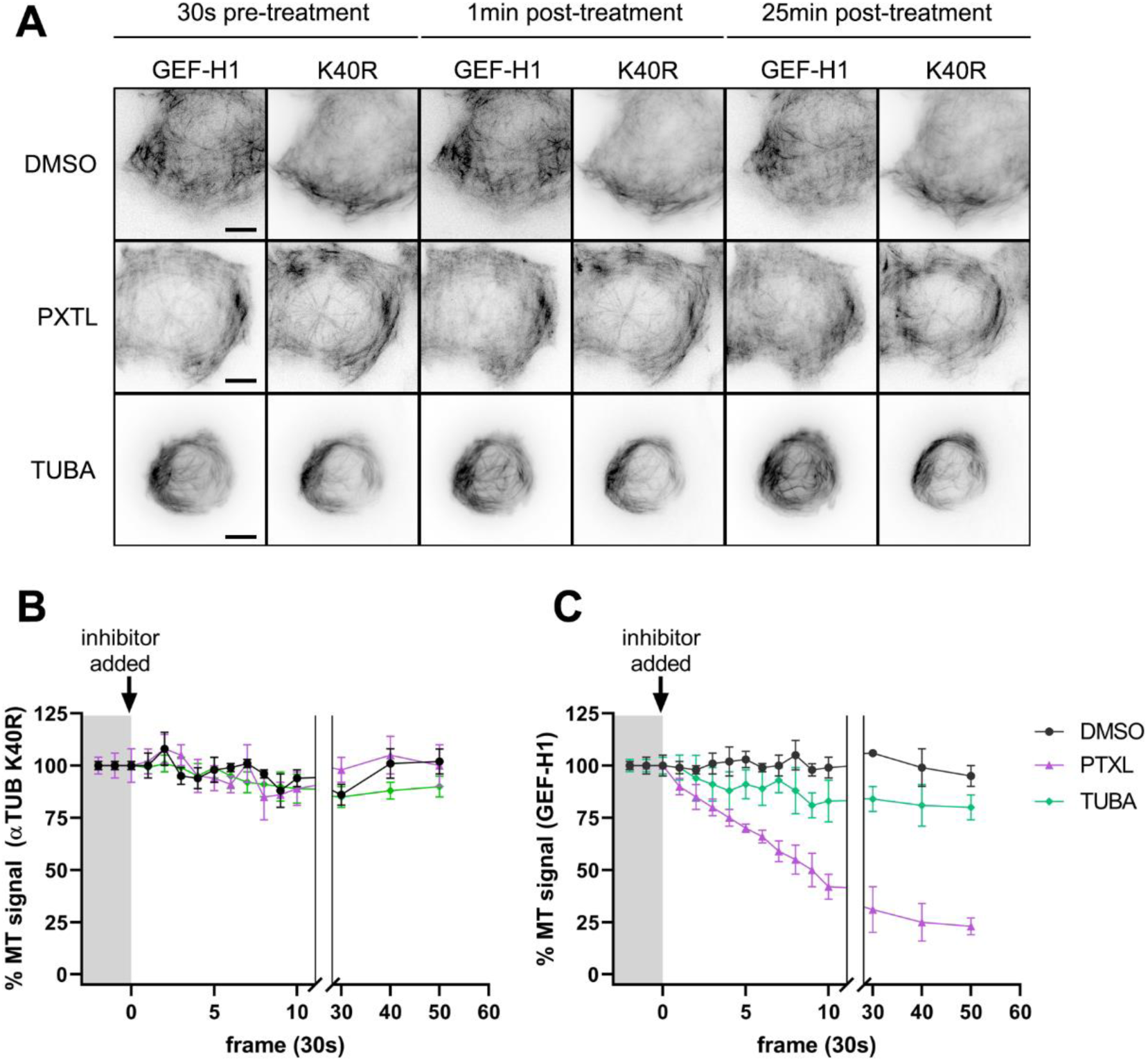
K40R mutation in α-tubulin partially inhibits PTXL-mediated release of GEF-H1 from microtubules. (A) Representative TIRF microscopy images from time-lapse movies of U2OS cells co-transfected with mCherry-α-tubulin K40R and GFP-GEF-H1. After 72 hours of co-transfection, cells were allowed to spread on fibronectin 2µg/mL coverslips. The next day, cells were placed under the microscope and treated with DMSO, 100 nM PTXL or 5 µM tubacin (TUBA) at time=0; then were imaged for thirty additional minutes. Left column represents a frame just before treatment, middle column 1 min post-treatment and right column 25 min post-treatment. Note the partial, slower dissociation of GEF-H1 from condensing microtubular structures in PTXL-treated K40R cells, in contrast to the complete, rapid release observed in WT microtubules (Fig. 7); and the absence of a statistically significant effect of tubacin on GEF-H1 localization in K40R cells. Bars = 20 µm. Images taken from supplementary videos 5-7. (B-C) Representative kinetics of MT signal of mCherry-α-tubulin K40R (B) and GFP-GEF-H1 (C). Quantification was performed as indicated in Material and Methods out of 12 independent movies per condition from three independent experiments.

### 2.4. PTXL-induced contractility is mediated by ROCK-NM2 and restrained by Arp2/3-dependent adhesion coupling

In addition to actomyosin contraction via ROCK-NM2, the GEF-H1-RhoA axis also controls other cellular processes, e.g. lamellipodial actin polymerization, particularly through the LIMK-cofilin pathway. To address whether altered actin polymerization contributes to the observed effects of taxanes on cellular contractility, we examined the specific effect of the treatments on lamellipodial actin organization in cells briefly stimulated with IGF-I to enhance protrusive activity **(Fig. 9)**. Vehicle (DMSO)-treated cells formed thin lamellipodia followed by a region sparsely enriched in actin bundles **(Fig. 9A, a)**. PTXL-treated cells displayed a similar organizational structure, but the lamellar region contained thicker, coalescing bundles and actin arches, consistent with increased NM2-mediated contractility **(Fig. 9A, b, white arrowheads)**. Tubacin-treated cells displayed an actin organization similar to that of vehicle-treated cells, but with overall thicker actin bundles behind the protrusion **(Fig. 9A, c, white arrows)**. In contrast, CK-666 (Arp2/3 inhibitor) and BMS-5 (LIMKi) induced collapsed lamellipodia **(Fig. 9A, d, g, blue arrowheads)**, confirming that both Arp2/3-dependent branching and LIMK-dependent cofilin inhibition contribute to lamellipodial maintenance in these cells. Interestingly, co-treatment with CK-666 and either PTXL or tubacin caused dramatic cell retraction, loss of actin fibers, and the emergence of actin-rich blebs characteristic of hypercontracted cells, with retraction and blebbing appearing more pronounced upon CK-666/PTXL co-treatment than CK-666/tubacin co-treatment **(Fig. 9A, e-f, blue arrows)**.

**Figure 9.**
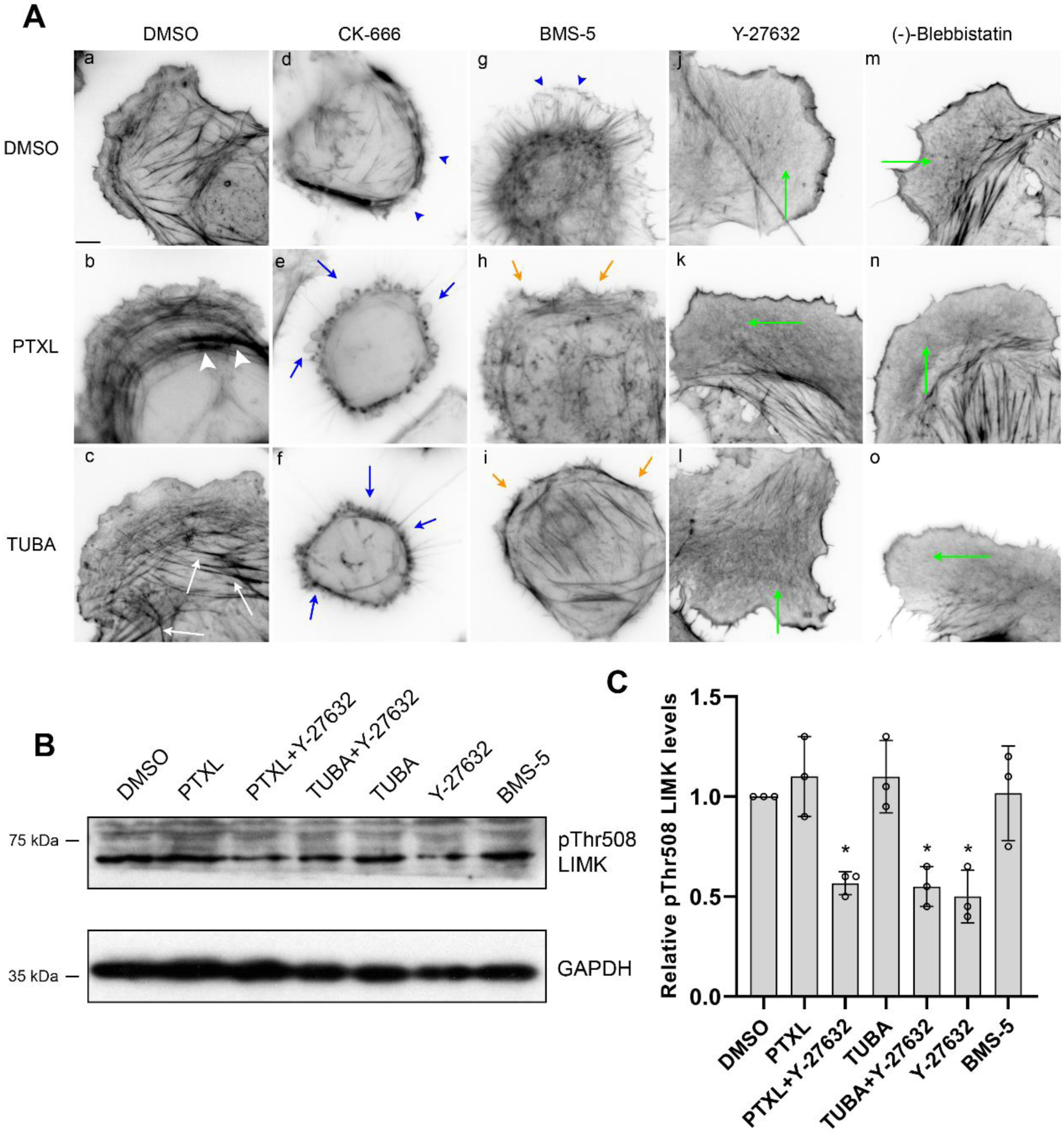
PTXL and tubacin increase lamellar actomyosin bundling through ROCK/NM2 independently of LIMK activation. (A) U2OS cells were allowed to spread overnight on fibronectin-coated dishes. The next day, cells were treated with DMSO (a, vehicle), PTXL (b, 100 nM) or tubacin (c, 5 µM) combined or not with Arp2/3 inhibitor CK-666 (d-f, 200 µM), LIMK inhibitor BMS-5 (g-i, 10 µM), ROCK inhibitor Y-27632 (j-l, 20 µM) or NM2 ATPase inhibitor (-)-blebbistatin (m-o, 30 µM). Treatment with tubacin was 2.5h. Treatment with the rest of the inhibitors was 30 min. To increase protrusiveness, cells were incubated with 100 ng/mL IGF-I for the last 15 min of the incubation. After that time, cells were fixed and stained for F-actin with AlexaFluor488-conjugated phalloidin. Images shown are representative of each condition. Experiment was performed three times. Scale bar= 10 µm. (B) U2OS cells were incubated with PTXL, tubacin, Y-27632, BMS-5, or PTXL or tubacin combined with Y-27632, using the same treatment times as in (A) (2.5h for tubacin, 30 min for the other inhibitors), lysed and blotted for phospho-Thr508 LIMK and GAPDH as loading control. A representative experiment of three performed is shown. (C) Quantification of LIMK phosphorylation in U2OS cells as in (A). Data are the mean ± SD of three independent experiments. Significance was determined with respect to DMSO-treated cells using Mann-Whitney U test. * p<0.05.

This is consistent with PTXL acting as a stronger inducer of contractility in our system. This suggests that Arp2/3-dependent dendritic actin polymerization normally counterbalances the additional contractile tension generated by microtubule-targeting drugs, preventing hypercontraction; loss of this protrusive counterforce (triggered by CK-666-induced Arp2/3 inhibition) leaves contractile tension unopposed, driving the cell into a hypercontracted, blebbing state. Notably, despite also disrupting lamellipodial architecture, LIMK inhibition did not phenocopy the dramatic retraction and blebbing observed with Arp2/3 inhibition **(Fig. 9A, h-i, orange arrows)**, suggesting that loss of lamellipodial actin organization per se is not sufficient to unleash hypercontraction. Instead, these results point to a specific role for Arp2/3-dependent branched actin in counterbalancing microtubule-induced contractile tension, potentially through its coupling to nascent adhesions and the substrate (35, 36), whereas LIMK/cofilin activity may primarily regulate actin filament turnover rather than substrate-coupled force transmission. Finally, co-treatment with the ROCK inhibitor Y-27632 and NM2 inhibitor blebbistatin negated the actin bundling effect of both PTXL and tubacin **(Fig. 9A, k-l, n-o)**. The effect was particularly prominent in the lamella, where bundled actin was largely absent (green arrows). Conversely, the rear part of the cell still displayed actin bundles, likely due to the increased stability of these structures, which endows them with partial resistance to the inhibitors.

In addition, we found that LIMK phosphorylation at T508, used as a readout of LIMK activation (37), was not induced by treatment with PTXL or tubacin, but was partially inhibited by Y-27632, a specific ROCK inhibitor **(Fig. 9B-C)**. Together, these experiments indicate that, although protrusion and contraction are interconnected processes, microtubule-targeting drugs increase contractility specifically through a ROCK/NM2-dependent mechanism. Arp2/3-dependent branched actin polymerization counterbalances this contractile tension, likely through its mechanical coupling to adhesions. LIMK-dependent actin turnover, in contrast, primarily shapes lamellipodial architecture without contributing to this force-buffering role.

### 2.5. PTXL does not increase the association of NM2-A with UNC45a

Increased cellular contractility can in principle arise from two distinct mechanisms: an elevated activity of the existing pool of NM2 filaments, which would be triggered by the GEF-H1/RhoA/ROCK axis described above, or an increase in the total number of NM2 filaments. PTXL stabilizes and straightens the microtubule lattice, and a recent study has suggested that the chaperone UNC45a binds curved, but not straight, microtubules (38), with microtubule straightening promoting UNC45a dissociation (39). Based on these findings, we asked whether PTXL-induced straightening could release UNC45a to drive new NM2-A filament assembly (24), analogous to the mechanism reported for *MYH9*-RD-causing NM2-A variants (40). To address this question, we initially assessed the association of UNC45a with NM2-A in tubacin- or PTXL-treated cells **(Fig. 10A)**. We found that neither PTXL nor tubacin increased the interaction of UNC45a and NM2-A **(Fig. 10A-B)**. Furthermore, examination of the subcellular localization of UNC45a, NM2-A and microtubules in control, PTXL- and tubacin-treated cells using TIRF microscopy revealed that UNC45a formed mostly linear filaments following the pattern of NM2-A. Conversely, we did not observe any patterning of UNC45a matching α-tubulin or acetylated microtubules **(Fig. 10C-D)**. We observed that UNC45a appeared brighter in filaments in which NM2-A was less prominent, whereas filaments and stacks brightly decorated with NM2-A were dimmer for UNC45a **(Fig. 10E)**. In filaments containing less NM2-A, UNC45a intercalated between existing NM2-A filaments and formed bright spots adjacent to them **(Fig. 10E)**. This suggests that UNC45a localization to filaments is important during their maturation but plays a less prominent role once NM2-A filaments become consolidated. Together, these data indicate that PTXL-induced microtubule straightening does not translate into increased UNC45a-dependent NM2-A filament assembly, ruling out an increase in total filament number as the source of enhanced contractility. Instead, the data point to increased activity of the existing filament pool, via the GEF-H1/RhoA/ROCK axis, as the main operative mechanism.

**Figure 10.**
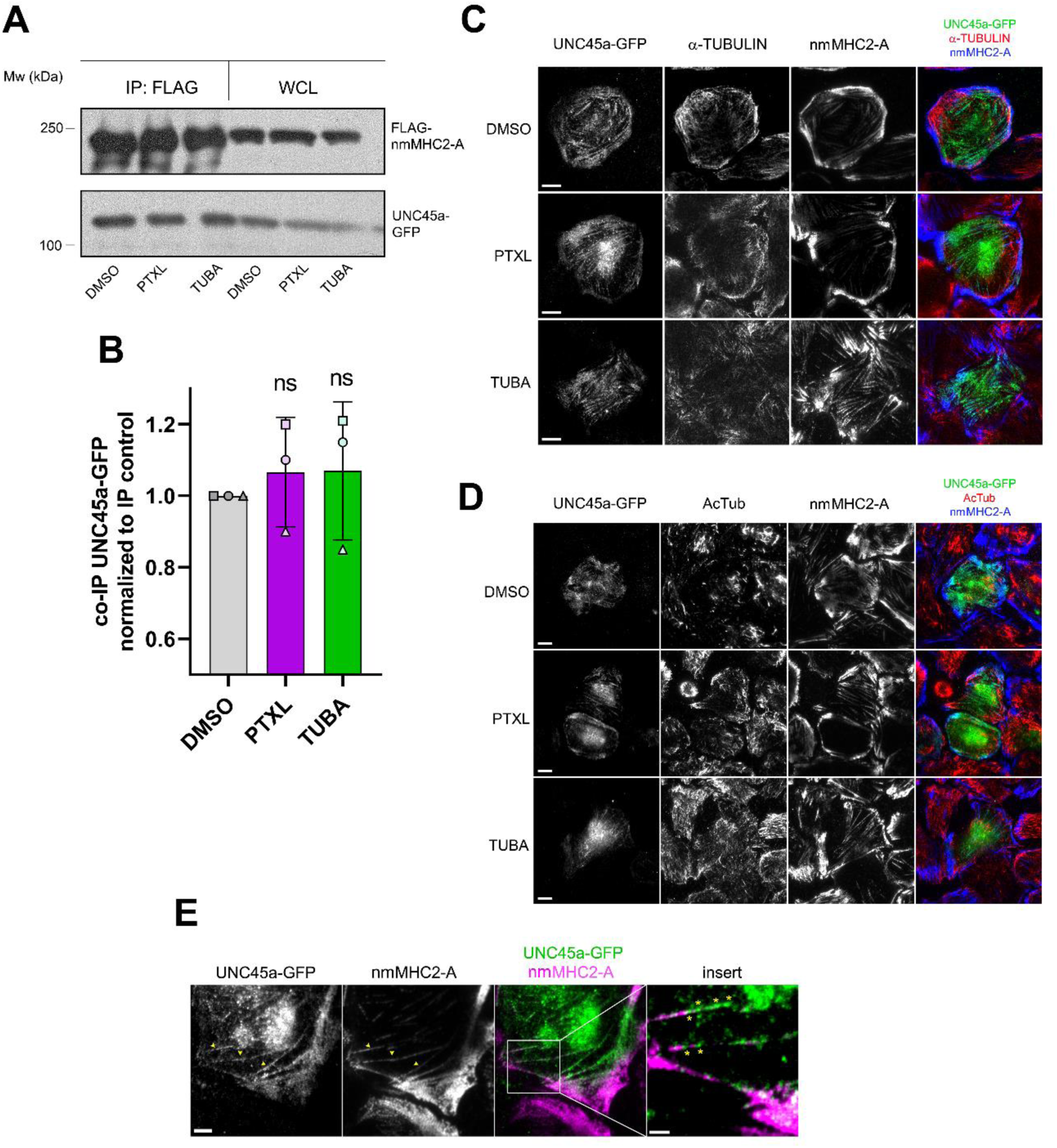
PTXL does not promote the association of UNC45a to NM2-A. (A-B) U2OS cells were co-transfected with FLAG-nmMHC2-A and UNC45a-GFP, treated as indicated with PTXL or tubacin, lysed and immunoprecipitated with anti-FLAG magnetic beads. Immunoprecipitates were separated by SDS/PAGE and blotted for FLAG and GFP as indicated. WCL, whole-cell lysates. Experiment is representative of three performed, and quantification of those experiments is shown in (B). ns, not significant differences (Mann-Whitney U test). (C-D) Representative TIRF microscopy images from U2OS cells transfected with UNC45a-GFP and treated as indicated. Cells were then stained for total α-tubulin (C) and acetylated tubulin (D). Scale bar=10 µm. (E) Representative TIRF image of U2OS cells transfected with UNC45a-GFP. Images illustrate that UNC45a clearly decorates posterior, thin filaments that contain few NM2-A molecules (arrowheads), whereas it is less represented in regions where NM2-A is highly prominent in bright stacks. Scale bar=5 µm. Insert illustrates the localization of UNC45a in filaments that contain few NM2-A molecules, forming bright spots between or adjacent to assembled NM2-A filaments (asterisks). Scale bar=5 µm.

### 2.6. NM2 is independently activated by PTXL and HDAC6 inhibition or αTAT1 depletion

Having shown that acute, PTXL-induced GEF-H1 release depends on K40 acetylation, we next asked whether basal, homeostatic levels of tubulin acetylation similarly shape steady-state actomyosin contractility, independent of any acute pharmacological trigger. We observed that inhibition of tubulin deacetylase HDAC6 using tubacin produced a significant accumulation of acetylated microtubules (41) and phosphorylation of RLC at Ser19 **(Fig. 1D-G)** concomitant with an increase in F-actin filament formation **(Fig. 11A-B)**. After 3h, this correlated with a modest, yet significant, increase in focal adhesion size **(Fig. 11C-D)**; adhesion number, which reflects a distinct process (adhesion assembly rather than maturation, as described (29, 36)), was not assessed in this experiment. Genetic depletion of HDAC6 with two independent shRNA sequences also increased tubulin acetylation and Ser19 RLC phosphorylation modestly **(Fig. 11E)**. Consistent with published results, HDAC6 removal modestly increased adhesion maturation **(Fig. 11F, H-I)**. While both sequences promoted adhesion maturation, the effect was statistically significant for one of them (labelled pLKO-HDAC6 HD), while the second sequence (pLKO-HDAC6 #39) showed a consistent trend in the same direction that did not reach significance (n=3 experiments). Consistent with a specific requirement for HDAC6 activity, tubacin failed to further increase focal adhesion maturation in HDAC6-depleted cells **(Fig. 11G-I)**, suggesting that both perturbations converge on a shared, rate-limiting step, and arguing against an off-target explanation for the phenotype. Together, these data indicate that both pharmacological and genetic HDAC6 loss increase actomyosin contractility, consistent with a lagged, acetylation-dependent effect on the actomyosin cytoskeleton (21). We next hypothesized that αTAT1 depletion would reduce RLC phosphorylation by enhancing GEF-H1 retention in microtubules. To test this hypothesis, we depleted αTAT1 using shRNA. Similar to cells depleted of αTAT1 using CRISPR/Cas9 gene editing (42), αTAT1 shRNA-transfected cells displayed significantly reduced acetylated tubulin (AcTub) **(Fig. 12A, light yellow and mauve arrows)** compared to non-transfected cells **(Fig. 12A, red arrow)**. Unexpectedly, we observed that AcTub-deficient cells did not display decreased RLC phosphorylation. Instead, they exhibited prominent pSer19 RLC phosphorylation in lamellae and reduced anteroposterior polarity, which is an indicator of increased isotropic contraction **(Fig. 12B-C).** These data indicate that the homeostatic levels of tubulin acetylation mediated by αTAT1 control RLC activation and the organization of actomyosin bundles. They also show that lowering the basal levels of acetylated tubulin redistributes tonic actomyosin contraction and subsequently controls the basal morphology of adherent cells in the post-spreading phase. In the context of the previous observations, this dataset further demonstrates that the activating effect of PTXL on NM2 is largely independent of homeostatic, αTAT1-dependent, microtubule acetylation.

**Figure 11.**
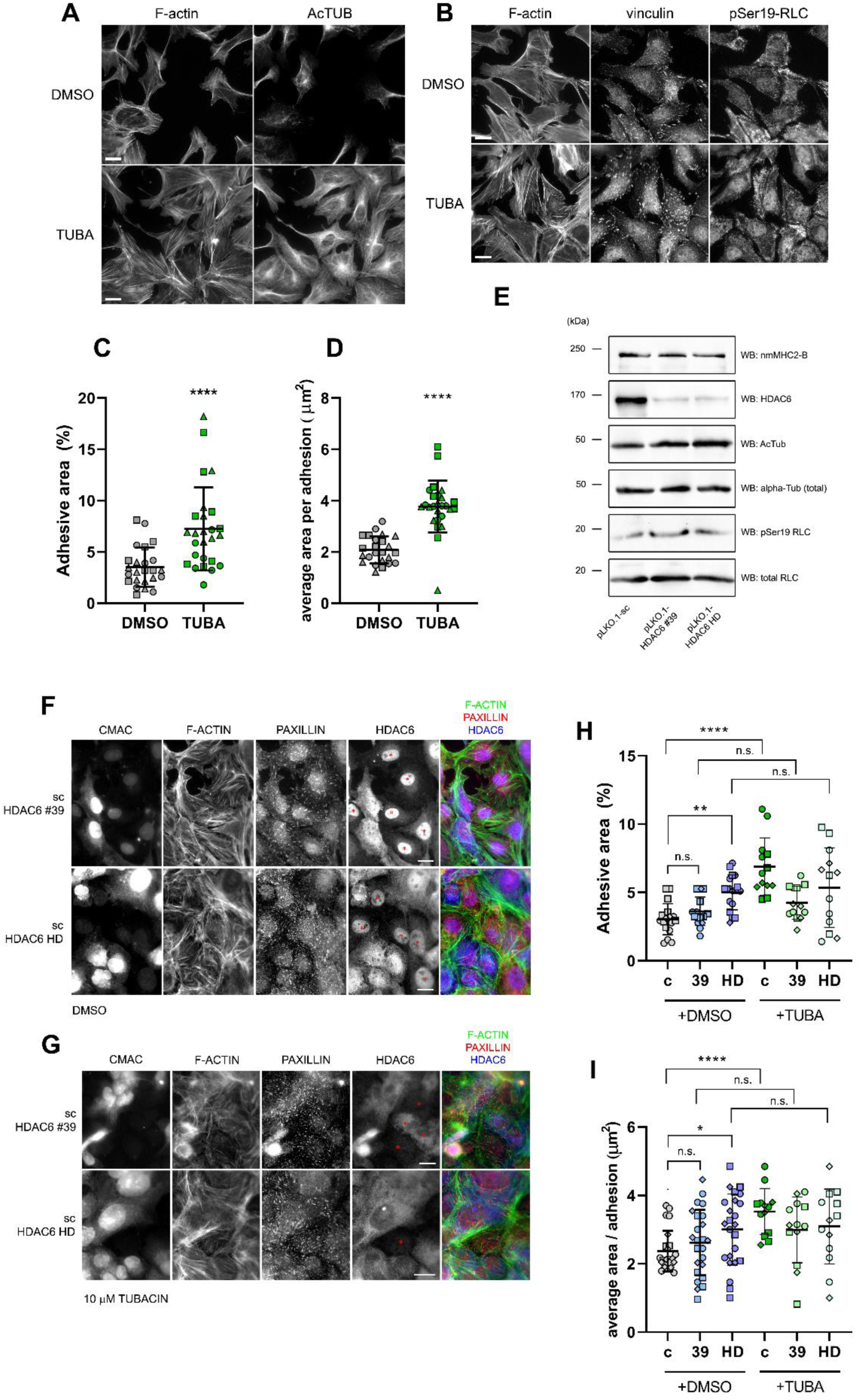
Pharmacological and genetic inhibition of HDAC6 increases contractility and focal adhesion maturation. (A) Representative fluorescence images of U2OS cells spread on fibronectin-coated coverslips treated with DMSO or 5 µM tubacin for 3h and stained for F-actin and acetylated tubulin (labelled AcTub). (B) Cells as in (A) were stained for F-actin, vinculin and pSer19 RLC. Scale bar= 10 µm. (C-D) Quantification of the percentage of adhesive area (C), representing the area of the cell positive for vinculin divided by the total area of the cell; (D) average area per adhesion, defined by vinculin staining as calculated in (Talayero and Vicente-Manzanares, 2020). Each data point represents a group of n≥20 cells from three independent experiments, each designated by a different symbol. (E) Representative Western blot analysis of control (scrambled) and HDAC6-depleted (shRNA #39 and HD) U2OS cells, showing efficient depletion of HDAC6 protein by both shRNA sequences, accompanied by a modest increase in acetylated α-tubulin and pSer19 RLC, despite no detectable change in NM2-B (nmMHC2-B) expression. Total α-tubulin and total RLC are shown as loading controls for AcTub and pSer19 RLC, respectively. (F) Representative fluorescence images of a mixture of DMSO-treated control (labelled CMAC) and HDAC6-depleted (not labelled) cells spread on fibronectin-coated coverslips. For clarity, HDAC6-depleted cells are denoted by red asterisks. Cells were stained as indicated with AlexaFluor488-phalloidin, paxillin and HDAC6. (G) Cells as in (F) were treated with 10 µM tubacin (instead of DMSO) for 3h and stained as indicated. (H-I) Quantification of adhesive area (H), representing the area of the cell positive for vinculin divided by the total area of the cell; (I) average area per adhesion, defined by vinculin staining as calculated in (Talayero and Vicente-Manzanares, 2020), of cells as in (F-G). Each data point represents a group of n≥20 cells from three independent experiments, each designated by a different symbol. Significances were determined using Mann-Whitney U test. ** p<0.01; *** p<0.001; **** p<0.0001.

**Figure 12.**
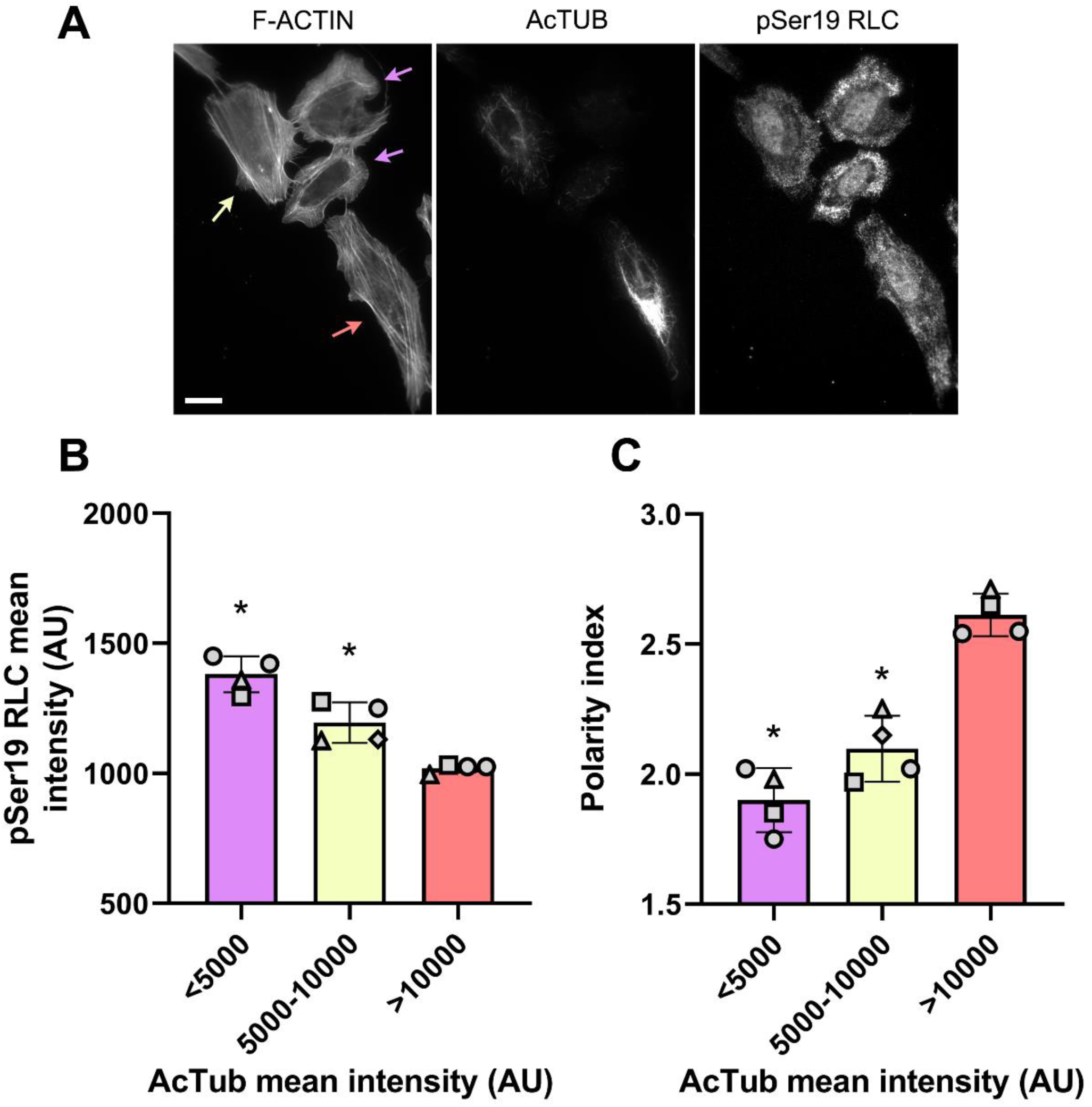
αTAT1 does not decrease pSer19 RLC levels but impairs anteroposterior polarity in U2OS cells. (A) Representative fluorescence images of U2OS cells transfected with a specific shRNA against human αTAT1. After 96 hours, cells were allowed to spread overnight on fibronectin-coated coverslips and stained for F-actin, acetylated tubulin (labelled AcTub) and pSer19 RLC. Colored arrows point to cells representative of each depletion level: red (high AcTub intensity, not depleted), light yellow (medium AcTub intensity, partial αTAT1 depletion), mauve (low AcTub intensity, deep αTAT1 depletion). Scale bar= 20 µm. (B-C) Quantification of pSer19 RLC (B) in arbitrary units and polarity index (C) with respect to grouped levels of AcTub (X-axis, AU). Each point represents >50 cells from four independent experiments. * p<0.05 (vs. the >10,000 AU (non-depleted) group, Mann-Whitney U test).

## 3. Discussion

Microtubule poisons remain a much-used tool in chemotherapeutic regimes to treat highly invasive forms of cancer (43). Of these, taxanes are the most commonly used, particularly for ovarian cancer and metastatic lung and breast cancer (44–46). While previous work had described the ability of microtubule polymerization inhibitors to increase cellular contraction via RhoA/NM2 (5, 17, 47), such observations were not consistent in the case of taxanes. For example, one study reported increased active RhoA in response to taxanes (48), whereas another reported that PTXL decreases collagen fiber contraction, which is inextricably dependent on cellular contractility (49).

Our experiments indicate that PTXL induces RLC phosphorylation and NM2 activation similar to nocodazole or VBL, suggesting a common mechanism of downstream activation. Our data also confirmed that the same mediators, GEF-H1 and RhoA, are involved in PTXL-induced NM2 activation and cellular contraction. A common assumption in the field is that microtubule disassembly releases GEF-H1, which then activates RhoA, indicating that GEF-H1 association to microtubules is a form of regulatory compartmentalization (50). Indeed, ARHGEF2/GEF-H1 is an active RhoGEF (51); thus limiting its intrinsic activity is essential to control the contractile homeostasis of the cell. Interestingly, PTXL does not disassemble the microtubule lattice, but condenses it (15). However, our data clearly show that PTXL quickly jettisons GEF-H1 from the microtubule network **(Figs. 6 and 7)**. This indicates that the binding of PTXL to the microtubule lowers its affinity for GEF-H1. A recent study proposed that the mechanosensitive acetylation of microtubules releases GEF-H1 from microtubules, thereby promoting actomyosin contractility (21). This is further supported by work using OptoTAT, an optogenetic tool that increases microtubule acetylation via αTAT1 (52), to elevate acetylated tubulin. We also observed a progressive decrease in GEF-H1 association when microtubules become acetylated in response to the HDAC6 inhibitor tubacin. Conversely, PTXL induces a near-instantaneous dissociation of GEF-H1 from microtubules, much faster than the effect of acetylation. Our data instead point to a mechanism by which microtubule stiffening by PTXL triggers the decrease in affinity for GEF-H1 independent of its effect on tubulin acetylation. In this regard, PTXL binds to the luminal region of the β-subunit of tubulin (53). It is likely that PTXL binding to the MT allosterically remodels the tubulin interdimer interface (54), resulting in longitudinal MT expansion (55). Recent work from the Steinmetz group has shown that the C1 domain of GEF-H1 binds both intra-tubulin dimer and inter-protofilament contacts on the outer surface of the microtubule (56), well away from the PTXL binding site (53). However, structural data explicitly support that the interaction of GEF-H1 with MTs is very sensitive to mechanical modifications to the microtubule, including changes in stiffness, curvature or bending (56). Based on these data, a likely model is that PTXL binding-dependent conformational increase of the microtubule lattice rigidity, which is distinct and complementary to that induced by GTP binding during polymerization (15), immediately lowers the microtubule lattice’s affinity for GEF-H1, resulting in its immediate release and downstream activation of actomyosin contractility.

Two recent studies have shed light on the specifics of microtubule stabilization by taxanes. In one, the authors describe that occupancy of the taxane binding site does not affect the straightness or stability of the tubulin protofilament but triggers its longitudinal expansion (53). The other identifies differences between the mechanism of action of PTXL (which expands the lattice, as reported above) and doublecortin, a protein that triggers microtubule compaction (55). Since expansion and compaction control PTM regulators of microtubules such as the tubulin detyrosinase VASH (57) and αTAT1 (58), it is likely that compaction/expansion controls GEF-H1 binding to microtubules.

Despite the strong experimental evidence that PTXL induces the rapid dissociation of GEF-H1 from microtubules through a mechanism involving microtubule rigidity and/or expansion, the observation that K40R α-tubulin significantly impairs, but does not completely abolish, PTXL-induced GEF-H1 dissociation warrants mechanistic consideration. K40 is the primary acetylation site of α-tubulin located in the lumen of the microtubule where it is accessible to αTAT1 (23, 24). Acetylation of K40 neutralizes the positive charge of the lysine ε-amino group, a modification that is absent in the K40R mutant. Importantly, while arginine also bears a positive charge at physiological pH, it cannot be acetylated, and therefore permanently mimics the unacetylated state of K40 regardless of αTAT1 activity. This distinction argues against a simple steric interpretation of the K40R phenotype and instead points to the electrostatic consequence of acetylation (that is, charge neutralization) as a key determinant of GEF-H1 dissociation efficiency.

Based on these observations, we propose a model in which PTXL-induced GEF-H1 dissociation involves two complementary and partially independent components. The first is mechanical: PTXL binding to the luminal face of β-tubulin triggers longitudinal lattice expansion (54), which is transmitted allosterically to the outer surface of the microtubule where GEF-H1 contacts the inter-protofilament and intra-dimer interfaces (36). This mechanical component alone is sufficient for partial GEF-H1 release, as evidenced by the residual dissociation observed from K40R microtubules. The second component is electrostatic: acetylation-dependent charge neutralization at K40 in the lumen facilitates the conformational propagation of the PTXL-induced expansion to the outer surface, enabling the rapid and complete dissociation of GEF-H1 observed from wild-type microtubules. In this model, the two components act synergistically: mechanical expansion provides the driving force, while electrostatic remodeling of the lumen tunes the efficiency and speed of allosteric transmission to the GEF-H1 binding site.

This framework also provides a plausible explanation for the differential kinetics of GEF-H1 dissociation induced by PTXL versus tubacin. PTXL simultaneously delivers both components: lattice expansion and, over time, increased acetylation, resulting in rapid and complete GEF-H1 release. Tubacin, by contrast, promotes acetylation without inducing lattice expansion, providing only the electrostatic component and producing a slower, more modest dissociation. The requirement for both components may represent a regulatory safeguard, ensuring that GEF-H1 is not released accidentally by either mechanical or chemical perturbation of the microtubule lattice alone, thereby preventing an inappropriate activation of the RhoA/ROCK/NM2 axis.

Our experiments also revealed that GEF-H1 compartmentalization is part of a homeostatic mechanism, acting in conjunction with protrusion-adhesion at the lamellipodium, that prevents hypercontraction. As shown here, taxanes drive contraction but do not affect the ability of the cell to undergo protrusion. As a result, adhesions are reinforced (they grow in size) due to the increased contractility. Alteration of the adhesion-protrusion coupling in these conditions, e.g. by Arp2/3 inhibition, drives hypercontraction and blebbing. One interpretation of these data is that, in these conditions, cells cannot “invest” the additional contraction in growing focal adhesions, retracting violently in response to increased centripetal contraction of the submembranous actin cortex. In this regard, Arp2/3 inhibition weakens the subcortical actin network, which has been linked to blebbing (59, 60). Our data concur with a previous report showing that abrogation of the transmission of the GEF-H1-dependent increase in contractility, e.g. by ROCK inhibition, suppresses membrane blebbing and restores lamellipodial polarity (61). Finally, elevated RhoA signaling also increases ERM-mediated membrane attachment first and NM2-dependent contractility second, a sequence that drives retractions and supports blebbing states (62).

Strikingly, our data also show that αTAT1 depletion alters the distribution of phosphorylated RLC, increasing its lamellar localization and decreasing post-spreading anteroposterior polarization. This was unexpected, particularly in light of the data indicating that microtubule acetylation promotes GEF-H1 release from the microtubule lattice (21, 52). Accordingly, αTAT1 depletion decreases K40 α-tubulin acetylation (22, 23), which should increase retention of GEF-H1 in microtubules. Our data thus suggest that the effect of αTAT1 depletion on actomyosin contractility may go beyond its role in tubulin acetylation/deacetylation, and that HDAC6 and αTAT1 may each control contractility, at least in part, through a tubulin acetylation-independent mechanism. Consistent with this, treatment with the HDAC6 inhibitor tubacin and genetic depletion of HDAC6 both increased focal adhesion assembly, in agreement with previous reports ((33), and this report). Inhibition as well as genetic depletion of HDAC6 does not preclude the involvement of other deacetylation substrates, such as cortactin or Hsp90 (63, 64), although the lack of effect of tubacin on GEF-H1 dissociation from K40R microtubules strongly argues that tubulin is the major substrate of HDAC6 involved in this effect. In addition, we showed that HDAC6 has catalytic-dependent and -independent functions (65). Likewise, αTAT1 controls microtubule dynamics independent of its tubulin acetylation activity, as a mutant unable to catalyze acetylation still promotes microtubule destabilization (66). The latter observation supports a model in which αTAT1 depletion promotes the generation of stable microtubules by alternative means, e.g. detyrosination, triggering the release of GEF-H1 from the microtubule lattice.

The ability of PTXL to promote actomyosin contraction suggests that enhanced contraction may control the development of PTXL-resistant cells during cancer therapy. In this regard, increased contraction in vivo may lead to the acquisition of additional migratory capabilities through mesenchymal-amoeboid transitions (67). This raises the possibility that resistant cells exposed to PTXL could become more migratory *in vivo*, paving the way for the use of anti-contractile adjuvant therapies. A recent example is the use of blebbistatin analogs to treat GBM in a preclinical model (68).

## 4. Material and Methods

### 4.1. Plasmids

EGFP-GEF-H1 was a gift from Alexander Bershadsky and has been described previously (69). UNC45a-EGFP was a gift from Pekka Lappalainen and was described elsewhere (24). pSUPER-GEF-H1 (human) contains the shDNA targeting sequence 5’-CAACATTGCTGGACATTTC -3’, which has been described elsewhere (31). pSUPER-RhoA (human) contains the shDNA sequence 5’-CAGTTCTGTGGTTTCATGT-3’, adapted from Ambion library. mCherry α-tubulin was obtained by direct subcloning of the mCherry gene into GFP-α-tubulin (originally obtained from Clontech, cat. no. 6117-1) using NheI/BsrGI restriction sites. mCherry α-tubulin K40R was obtained by direct mutagenesis of the K40 codon using GeneTailor kit (Invitrogen). Primers were: 5’-ATGGCCAGATGCCAAGTGACA**ga**ACCATTGGGGG-3’ and 5’-GTCACTTGGCATCTGGCCATCGGGCTGGAT-3’. pSUPER-ATAT1 (human) contains the shDNA sequence 5’-CTGGATGATCGTGAGGCTC-3’, adapted from Ambion library AM16704. pLKO-HDAC6 #39 sequence was obtained from Merck Mission shRNA library (cat. no. TRCN0000004839). pLKO-HDAC6 HD contains the sequence 5’-GCTGCACCGTGAGAGTTCCAACTTT-3’ as described in (70), and was cloned as a shDNA into pLKO.1 following standard practice (https://www.addgene.org/protocols/plko/).

### 4.2. Antibodies and reagents

Antibodies against the following antigens were used: acetylated tubulin (RRID:AB_609894), tyrosinated tubulin (RRID:AB_2890657), total RLC (clone MY-21, RRID:AB_477192), vinculin (clone hVin-1, RRID:AB_477629) and FLAG (clone M2, RRID:AB_439685) were from Merck; GAPDH (RRID:AB_10613283), nmMHC2-A (RRID:AB_2734686) and nmMHC2-B (RRID:AB_2749903) were from Biolegend; phospho-RLC (Ser19) rabbit polyclonal antibody was from Rockland Immunochemicals (RRID:AB_2610739) or from Cell Signaling Technology (RRID:AB_330248), as indicated in the figure legends; pThr18/pSer19 RLC (RRID:AB_2147464) was from Cell Signaling Technology; GEF-H1 (RRID:AB_2818944) was from Abcam; RhoA (RRID:AB_628218) and GFP (RRID:AB_627695) were from Santa Cruz Biotechnology; HDAC6 (RRID:AB_10597094) was from Proteintech; paxillin (RRID:AB_397952) was from BD Biosciences. Highly cross-linked anti-mouse (H+L) and anti-rabbit secondary antibodies coupled to AlexaFluor488, 568 or 647 (RRID:AB_2534069; -11004/, RRID:AB_2534072; RRID:AB_2633277; RRID:AB_143165; RRID:AB_143157 and RRID:AB_2535812), and AlexaFluor647-Phalloidin (A30107) were from Thermo. Fibronectin (F-2006) was from Merck. Paclitaxel, vinblastine and tubacin were from MedChemExpress. IGF-I (PHG0071) was from Thermo.

### 4.3. Cell lines

U2OS (RRID:CVCL_0042), OVCAR-8 (RRID:CVCL_1629), SK-OV-3 (RRID:CVCL_0532) and NIH-3T3 (RRID:CVCL_0594) cells were maintained in DMEM medium supplemented with 10% FBS, 100 U/mL penicillin/streptomycin and 1% non-essential amino acids. Cells were routinely sub-cultured when 80% confluent and used for a maximum of 20 passages.

### 4.4. Microtubule fractionation, Western blot and quantification

Cells were lysed using 1% NP40 in TBS (Tris-buffered saline) containing 10 μM ATP, 10 mM MgCl_2_ and 1 mM EDTA, plus protease and phosphatase inhibitors (10 μM aprotinin, leupeptin and 1 mM PMSF; and 1 mM NaF, Na_3_VO_4_, Na_4_P2O_7_, all from Merck), clarified by centrifugation (17000×g, 15min, 4°C) and quantified with a BCA Protein Assay Kit (Pierce, part of Thermo Scientific). Samples were mixed with an equal volume of 2× SDS protein sample buffer containing 40% glycerol, 240 mM Tris/HCl pH 6.8, 8% SDS, 0.04% bromophenol blue and 5% beta-mercaptoethanol. For microtubule fractionation, 1.5×10^6^ cells in p60 dishes were treated with the anti-mitotic drugs for 3 hours. After the treatment, cells were scraped using PEM Buffer (PIPES 100mM pH6.9, EGTA 10mM, MgSO_4_ 1mM, 3M Sucrose, 1% Triton X-100 1% and protease/phosphatase inhibitors). Where indicated, the lysis buffer was supplemented with 100 nM VBL or 100 nM PTXL. Cell lysates then were centrifuged at 5000 ×g for 10 mins at 37°C to separate the pellet (tubulin polymers) and the supernatant (soluble tubulin dimers/monomers). Insoluble pellets were mixed with 200 μL 2× Laemmli buffer, supernatants were mixed 1:1 with 4× Laemmli buffer, and they were boiled for 5 mins at 95°C. In all cases, proteins were resolved by SDS-PAGE (sodium dodecyl sulfate polyacrylamide gel electrophoresis) and transferred to Immobilon^®^-FL PVDF 0.45 µm pore size membranes (Merck-Millipore) in a Bio-Rad wet blotting system. Membranes were blocked using TBS containing 5% BSA for one hour under continuous stirring. Primary antibodies were incubated in TBS with 0.1% Tween-20 (TBS-T) overnight at 4°C in a shaker and washed three times 10 min each with TBS-T. Peroxidase-labelled secondary antibodies were incubated for one hour and rinsed again three times for 10 min each with TBS-T. Chemiluminescence was developed using Immobilon^®^ Forte Western HRP Substrate (Merck-Millipore), and images collected in a Classic E.O.S. photographic development system from AGFA and X-rays blue medical film (AGFA/Fujifilm). Films were scanned at high resolution (600dpi minimum) in a Canon 9000F mark II scanner, band intensity was analyzed with ImageJ (densitometry plugin) and results were normalized to the expression of the loading controls as indicated for each figure.

### 4.5. Immunofluorescence, imaging and focal adhesion quantification

Cells were transfected with the indicated plasmids and incubated for 48-72h (72-96h for knockdowns). Where indicated, control cells were labelled using 1 µM CMAC according to the manufacturer’s instructions. 10^5^ total cells were allowed to adhere to a #1.5H, 12mm round coverslip coated with 2 µg/mL human fibronectin for 3h. Cells were subsequently fixed with 4% paraformaldehyde (Pierce Thermo) in PBS for 10 min, rinsed three times in TBS and stained as indicated. Cells were observed in a Leica Thunder Tissue Analyzer equipped with a 63× objective (NA 1.45) and appropriate LED laser lines and filters for DAPI (used to image CMAC), EGFP, AlexaFluor568 and AlexaFluor647 imaging. Images (20-30 per condition in at least three independent experiments) were collected using LAS X and focal adhesions quantified as described (71), using vinculin or paxillin immunostaining interchangeably, as the validity of both markers for these measurements was reported in the same study. Briefly, we subtracted background, applied the CLAHE (Contrast Limited Adaptive Histogram Equalization) algorithm to create a high-contrast image that was subject to the threshold command of ImageJ and quantified using the Analyze Particles ImageJ built-in feature. % of adhesive area was obtained by dividing the adhesive area of the cell (determined as described above) by the total area of the cell (determined manually). Polarity index was calculated as described elsewhere (29). Briefly, we divided the length of the long, migration-defined axis by the perpendicular axis passing through the center of the nucleus of the cell. With this metric, the PI of a cell can only be ≥1, and a perfectly round cell has a PI=1.

### 4.6. TIRF microscopy

U2OS cells were co-transfected with GFP-GEF-H1 (0.8 µg/well) and α-tubulin-mCherry, WT or K40R (0.2 µg/well). After 72h, cells were allowed to adhere to a #1.5H, 25-mm round coverslip coated with 2 µg/mL human fibronectin for 30 min. Coverslips were attached to Attofluor^TM^ devices (Thermo) and the reservoir filled with 750 µL DMEM medium (no phenol red, Gibco cat. no. 11520556) supplemented with 10% FBS, 100 U/mL penicillin/streptomycin and 1% non-essential amino acids. Cells were imaged in a Leica inverted Thunder Tissue Analyzer equipped with a 63× objective (NA 1.45), appropriate LED laser lines and filters for EGFP and AlexaFluor568 imaging and an Infinity TIRF DMi8 S module (Leica). Cells were imaged under TIRF conditions with a 100-120 nm evanescent field. One image was captured every 30s, and movies are shown at 24 frames/sec. For multicolor samples **(Fig. 7A-B, E, 8A and 10C-E)**, evanescent field penetrance ranges and azimuth position were optimized for each wavelength individually, within the 90-130nm range (penetrance) and 0-180° (azimuth).

### 4.7. Co-immunoprecipitation of nmMHC2-A and UNC45a

U2OS cells (2× 10-cm diameter tissue culture dishes/condition at 60% confluence) were co-transfected with equimolar concentrations of plasmids encoding FLAG-nmMHC2-A and UNC45a-GFP (24). After 48h, cells were treated as indicated and resuspended in 1mL lysis/MgATP buffer. Lysates were incubated at 4°C for 15 min, clarified by centrifugation at 14000 ×g. Then, clarified lysates were incubated with 10 µL of either GFP-trap or FLAG-M2-magnetic beads for 120 min at 4°C under continuous stirring. Magnetic beads were rinsed three times with 1 mL lysis/MgATP buffer and resuspended in 100 µL 1× Laemmli buffer. The levels of the associated molecule (nmMHC2-A or UNC45a) were assessed by Western blot and densitometric quantification using ImageJ.

### 4.8. Statistics

In general, a normality test (Shapiro-Wilk) was first applied to the conditions under comparison. Significance of data deviating from a normal distribution was calculated using Mann-Whitney U test. Due to the importance of these comparisons for the interpretation of the data throughout the study, significance is indicated either in the figure itself or in its legend.

## DECLARATIONS

### Ethics approval and consent to participate

Does not apply.

### Consent for publication

The authors provide full consent for publication. There is no published or unpublished data requiring additional consent.

### Availability of data and material

Original data and unique plasmids and other materials will be made available by the authors upon reasonable request.

### Authors’ contributions

GAJ, data acquisition and analysis, initial draft preparation and manuscript correction; RPD, data acquisition; HRS, data acquisition; MGC, data acquisition; VCT, data analysis; MVM, project funding and management, global project design, data acquisition and analysis, manuscript writing and correction.

## Additional acknowledgements

The authors thank IBMCC (Salamanca) Advanced Cellular Imaging (Ana Isabel García de Vega) for technical support. The authors used Claude, an LLM developed by Anthropic, to assist with language editing of the manuscript; all scientific content, data interpretation, and conclusions are the sole responsibility of the authors.

## Authors’ institutional addresses

Molecular Mechanisms Program, Centro de Investigación del Cáncer/ Instituto de Biología Molecular y Celular del Cáncer, Consejo Superior de Investigaciones Científicas (CSIC) and University of Salamanca, 37007 Salamanca, Spain.

## Corresponding author

**Supplementary material.**

**Video 1.** Representative TIRF time-lapse movie of U2OS cells co-transfected with mCherry-α-tubulin (magenta) and GFP-GEF-H1 (green), treated with DMSO. Corresponds to Fig. 7A.

**Video 2.** Representative TIRF time-lapse movie of U2OS cells co-transfected with mCherry-α-tubulin (magenta) and GFP-GEF-H1 (green), treated with VBL. Corresponds to Fig. 7A.

**Video 3.** Representative TIRF time-lapse movie of U2OS cells co-transfected with mCherry-α-tubulin (magenta) and GFP-GEF-H1 (green), treated with PTXL. Corresponds to Fig. 7A.

**Video 4.** Representative TIRF time-lapse movie of U2OS cells co-transfected with mCherry-α-tubulin (magenta) and GFP-GEF-H1 (green), treated with TUBA. Corresponds to Fig. 7A.

**Video 5.** Representative TIRF time-lapse movie of U2OS cells co-transfected with mCherry-α-tubulin K40R (magenta) and GFP-GEF-H1 (green), treated with DMSO. Corresponds to Fig. 8A.

**Video 6.** Representative TIRF time-lapse movie of U2OS cells co-transfected with mCherry-α-tubulin K40R (magenta) and GFP-GEF-H1 (green), treated with PTXL. Corresponds to Fig. 8A.

**Video 7.** Representative TIRF time-lapse movie of U2OS cells co-transfected with mCherry-α-tubulin K40R (magenta) and GFP-GEF-H1 (green), treated with TUBA. Corresponds to Fig. 8A.

